# Spatiotemporal control of cortical centrin patterning by regionalized Sfi1 family scaffolding proteins in *Stentor coeruleus*

**DOI:** 10.1101/2025.08.11.669701

**Authors:** Connie Yan, Niklas Steube, Gautam Dey, Wallace F. Marshall

**Affiliations:** Department of Biochemistry & Biophysics University of California, San Francisco San Francisco, CA; Cell Biology and Biophysics European Molecular Biology Laboratory, Heidelberg, Heidelberg, Germany

**Keywords:** Stentor coeruleus, ciliate patterning, single-cell regeneration, oral regeneration, cellular contraction, expansion microscopy

## Abstract

Patterning is fundamental to the development and maintenance of organisms, ensuring functional and structural organization. While patterning is well studied at the level of multicellular organisms, even single cells need to undergo morphogenesis and form spatial patterns. *Stentor coeruleus* is a large ciliate that has been a classical system for studying patterning and morphogenesis due to its distinctive shape and organization of easily visible cortical structures which show a clear localization along the anterior-posterior body axis. The cortex of *Stentor* is made of two cytoskeletal layers, one composed of microtubules, and one composed of a network of centrin-family EF hand proteins, which form a branched network in the anterior half of the cell and long thick bundles known as myonemes in the posterior half. Sfi1 family proteins scaffold the assembly of centrin filaments throughout the eukaryotes, and function as scaffolding proteins in myonemal contractile systems found in some ciliates. A set of Sfi1 proteins upregulated during regeneration in *Stentor* were found to have a role in maintaining anterior/posterior differences in centrin patterning. Loss of this distinct anterior-posterior patterning leads to defects in regeneration and contraction. Using RNAi-mediated knockdown of Sfi1 genes, we have found that cells fail to regenerate their oral apparatus, with early expressed Sfi1 genes more critical for oral primordium (assembly of organized basal bodies that become the new oral apparatus) formation than those expressed later. Additionally, different Sfi1 proteins appear to be recruited sequentially to the growing primordium in an order that matches the time at which they are required for progression of regeneration. Knockdown of Sfi1 isoforms that results in reduced myoneme cables impairs contraction such that cells fail to contract in response to a stimulus or do not fully contract. These findings suggest a model in which regionalized differences in patterning of cytoskeletal assemblies can be modulated by regionalized localization of scaffolding proteins.

## Introduction

In the context of organismal development, patterning refers to the process by which certain biological structures and functions emerge in a predictable, organized way. Such developmental processes as axis formation, patterning and morphogenesis have been extensively studied on an organismal or tissue level, but much less so on a subcellular level. Yet cells have complex shapes and forms which are usually essential for their function, as well as intracellular patterning which ensures that components such as organelles are distributed correctly within the cell. Spatial and temporal organization of proteins within cells can also encode information about the cell’s geometry and developmental stage. But how do such patterns form within a single cell?

Ciliates provide an excellent opportunity for exploring cellular patterning, given their highly standardized cell shapes and readily observable surface structures that serve as clear landmarks for morphological processes^1,2^. The entire surface of these cells is lined with parallel rows of cilia that run along the anterior-posterior axis with defined spacing between adjacent rows. Additionally, in many ciliates, structures such as the gullet or the contractile vacuole occupy well-defined positions along both the anterior-posterior and circumferential axes. In some instances, the formation of a distinct structure or process on the cell’s surface is an expression of polarization or axiation. Polarization of the cytoskeleton plays a key role in providing a cell with positional information, for instance, actin filaments can guide the direction of movement in migrating cells, and the mitotic spindle helps position chromosomes in dividing cells^3^. Additionally, it has been shown that mRNA can be regionalized in ciliates and that microtubules may play a role in transport^4^. This spatial regulation of gene expression at the subcellular level highlights that even unicellular organisms can create and maintain internal asymmetries or gradients.

The giant ciliate *Stentor coeruleus* is a classical model system for studying regeneration, patterning and morphogenesis at a single cell level^5,6^. *Stentor coeruleus* is large, up to 1mm in length, with a highly complex body plan and a wide range of behaviors, including the ability to learn, respond to stimuli, and regenerate^7–9^. At its anterior is its oral apparatus, which consists of the membranellar band and mouth or gullet (Figure 1A). The membranellar band is composed of clusters of specialized cilia used for feeding. Along its body are pigmented blue stripes whose width varies in a stereotyped manner, decreasing from wide to narrow stripes as a function of longitudinal angle. The region where the narrowest and widest stripes meet functions as the oral primordium site during regeneration and cell division. Alternating in between these stripes are clear stripes lined with thousands of basal bodies, from which cilia extend, and which are also linked into parallel microtubule bundles that occupy the gaps between the pigment stripes^10,11^. Two fiber systems are contained in the cortical stripes, one a cytoskeletal microtubule-based fiber and the other, a contractile, myonemal system^6^ made of centrin proteins^12^.

**Figure 1.**
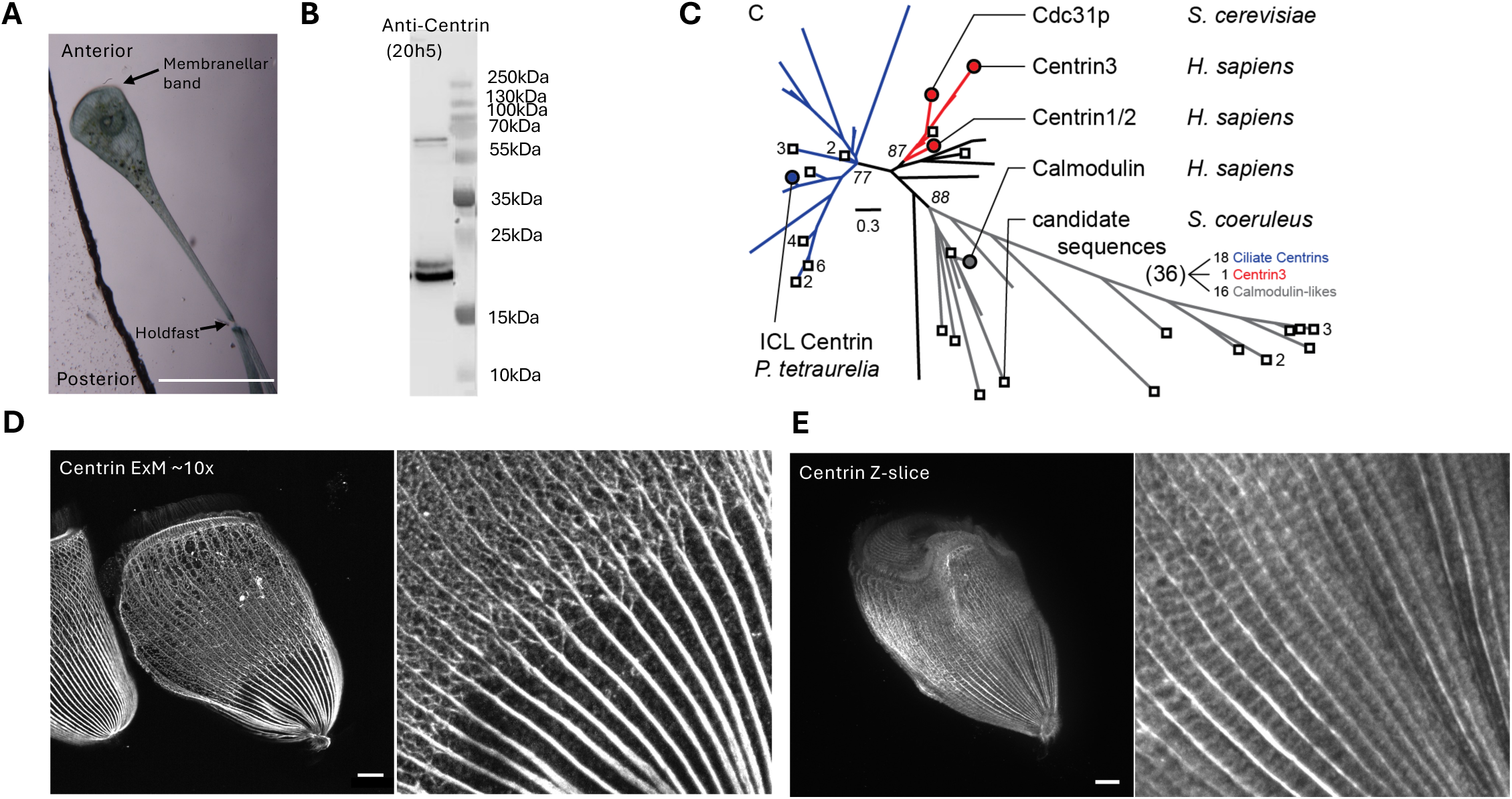
Regional differences in centrin patterning on *Stentor* cortex. (A) Brightfield image showing the general morphology of *Stentor* in its extended state, noting the key features of the anterior and posterior. The membranellar band, a feeding organelle consisting of clusters of specialized cilia sits at the anterior of the cell. The posterior stalk-like portion of the cell terminates at the holdfast, used for attaching to substrate. The stalk-like part of the cell can undergo rapid contraction. All following *Stentor* images are oriented with the membranellar band at the anterior and holdfast at the posterior. Scale bar 100um (B) Western blot of *Stentor* lysates using monoclonal anti-centrin (clone 20h5) antibody show bands at estimated centrin sizes (∼18-23kDa) and predicted larger calmodulins (55-70kDa) (C) Maximum likelihood tree of 36 *Stentor* sequences (squares) with *Paramecium tetraurelia*, *Saccharomyces cerevisiae* and *Homo sapiens* centrins as hallmarks. ICL, infracilliary lattice. Numbers count sequences near tips. Italic numbers are ultrafast bootstrap support values of 1000 replicates. Branch-lengths represent average amino acid substitutions per site. (D) Representative example of maximum intensity projection of confocal z-stacks showing *Stentor* centrin immunofluorescence on Magnify expanded cells. Close-up (right panel) highlights the stark change in pattern in the anterior and posterior of the cell and the intricate anterior centrin bundles in between the cortical rows. Scale bar 10um, adjusted for expansion. (E) A single confocal z-slice of centrin immunofluorescence, without expansion. Ladder “rungs” are seen in the posterior between the centrin containing cortical rows that run along the cell. These “rungs” are a distinct pattern from the anterior centrin networks and are distinct from body cilia which line the cell. Scale bar 10um.

Centrins are a highly conserved family of small EF-hand calcium binding proteins found across all eukaryotic lineages. They are primarily associated with fibrous structures at centrioles and basal bodies where they contribute to a diverse array of critical cellular functions, including centriole orientation, biogenesis and duplication, as well as nuclear export and DNA damage repair^13–18^. Beyond these canonical roles, centrins also participate in the organization of microtubule-based cytoskeletal elements, such as the mitotic spindle and motile cilia^19^, highlighting their diverse functional roles across both structural and regulatory contexts.

In contractile ciliates, centrins acquire an additional, specialized role in mediating calcium-dependent contraction. Previously, ultrastructural studies across multiple contractile ciliates have shown two morphologically distinct types of cortical fiber systems, both of which are thought to contribute to cellular contraction. One system is microtubule based, often associated with classical cytoskeletal functions like support and intracellular transport and the other consisting of fibrils or myonemes, which are thought to mediate rapid contraction in response to calcium influx^10,11,20,21^. These myonemal fibers contain centrin or related EF-hand family proteins and are regulated through calcium binding mechanisms, allowing for ATP independent, high-speed contraction. Centrin binds to Ca^2+^, undergoing a conformational change which allows cells to contract without ATP hydrolysis^22,23^. Contraction involves a dramatic change in morphology of the filaments in the myonemes, with the filaments of the myonemes coiling into helical coils during the contraction and becoming thicker and shorter^24,25^. A related fibrous structure that also contracts in response to changes in Ca^2+^ concentration is the vorticellid spasmoneme^26^. Its principal constituent, spasmin^23^, has homology to EF-hand domain proteins such as centrin^27^, and is similarly activated by calcium binding. In addition, the spasmoneme also has elastic properties that are not dependent on the Ca^2+^ concentration^26^. Taken together, these systems suggest that centrin containing fibrous assemblies are capable of integrating both active Ca^2+^ triggered contraction and passive elastic recoil. These properties of stimulus driven mechanical response and elasticity may be hallmark features of centrin-based cytoskeletal structures, enabling rapid and efficient cellular movements in ciliates and potentially other mechanosensitive eukaryotic cells.

The assembly of centrin into myonemes with defined orientations relative to the rest of the cell cortex raises the question of what positional information drives their patterned assembly. Centrin-binding partners are likely responsible for the specificity of localization and function of centrin. The founding member of a family of centrin-scaffolding proteins was identified in *Saccharomyces cerevisiae,* where centrin has been shown to have two clear roles in the spindle pole body (SPB): duplication of the SPB and as a constituent of the contractile fibers within and attached to the microtubule-organizing center, which are calcium sensitive^28^. A novel centrin binding protein was discovered in yeast, which co-localized with the centrin, Cdc31p, at the SPB half-bridge. This protein, Sfi1p, is a 120kDa polypeptide, containing 21 motifs with the consensus sequence AX_7_LLX_3_F/LX_2_WK/R, each binding to centrin at a 1:1 molar ratio^28,29^. Orthologs of this protein Sfi1p, were subsequently identified in many other diverse species.

Within the ciliates, sequence analysis has identified several large EF-hand binding proteins containing Sfi1 repeats, suggesting that this family of proteins plays a conserved and functionally diverse role across different species. In *Paramecium tetraurelia*, these proteins have been characterized and shown to be essential for the formation and maintenance of the infraciliary lattice (ICL), a calcium-responsive, contractile meshwork of fibers, containing Sfi1-centrin complexes, that is a prominent component of the *Paramecium* cytoskeleton. The ICL underlies the cell cortex and coordinates with basal bodies and other structural components to support cell shape, contractility, and mechanical stability ^30,31^. In *Tetrahymena*, Sfi1 homologs including sfr1 and sfr13 have been shown to play critical roles in basal body separation, spacing, and cortical organization. These proteins are required for the generation and stability of basal body rows, highlighting the importance of Sfi1-centrin interactions not only in contractile systems but also in templated organelle duplication and spatial patterning of the cell cortex^32,33^. Supporting this role, Sfi1 repeat proteins have also been identified in *Spirostomum minus*, where they functioning in ultra-fast Ca^2+^ mediated contraction^34,35^. In *Spirostomum*, the Sfi1 homologs, GSBP1 and GSBP2, are very large in size and assemble into fibrillar scaffolds that contract in response to calcium signaling, in coordination with their centrin family binding partner, spasmin^34^.

In *Stentor*, the myonemes or “M fibers”^36^ are described as discrete and prominent bundles in the posterior but linked by extensive side branches in the anterior^10,11,36^. These fibers are confirmed to be centrin containing where centrin is largely in the posterior cables of the cell with the fibers narrowing as they approach the anterior end of the cell^12^. At the anterior is a mesh-like network of centrin, creating a continuous sheet around the entire cell^36^, presumably to counter the change in internal pressure during rapid contraction where the stalk of the cell contracts to 20-25% of its full extended length^25^. The anterior end of *Stentor*, in contrast, undergoes very little extension or contraction. The frontal field, a region of the cortex at the anterior end of the cell surrounded by the membranellar band, is moderately contractile, but being circular in shape, contracts uniformly in all directions instead of along a single axis^37^. Perpendicular connections to the “M fibers” have been described^6,10,38^ although it was not known whether or not they contain centrin. How does the same centrin protein assemble into such distinct structures in different parts of the cell? Because of the association of Sfi1 family proteins with centrin in other contractile ciliates^30,32,34,35^, it is likely that Sfi1 is also involved in cortical fiber systems and contraction in *Stentor*. A prior proteomic analysis of dissected *Stentor* cells indicated that different Sfi1 family members were differentially localized along the anterior-posterior body axis^39^. Could the regionalized patterning difference in centrin into networks versus cables be due to regionalized localization of different Sfi1 proteins?

In order to investigate the interplay of Sfi1 family proteins and centrins in cortical patterning in *Stentor*, we have visualized the centrin based cytoskeleton using expansion microscopy and functionally analyzed a sub-set of Sfi1 proteins encoded by genes that are upregulated during oral regeneration. Our results indicate that the specific Sfi1 isoforms analyzed in this study co-localize with a specific subset of the centrin patterning, suggesting a selective spatial overlap rather than a uniform co-distribution. Furthermore, Sfi1 proteins exhibit distinct organization along the longitudinal axis of the cell, and knockdown of specific Sfi1 members led to altered spatial arrangement of centrin fibers, together pointing to a role in establishing or maintaining regionalized patterning of centrin within the cell cortex. Additionally, we find that Sfi1 proteins may have a functional role in oral regeneration. Specifically, different Sfi1 family members appear to be recruited sequentially to the growing oral primordium, supporting the idea that the primordium scaffold is assembled in a staged manner, with each Sfi1 isoform potentially scaffolding centrin at distinct phases of regeneration. Knockdowns of early expressed Sfi1 genes result in more severe regeneration defects, while late-expressed Sfi1 genes appear to be less essential, possibly due to functional redundancy or their role in later, less critical stages of oral primordium development. Given its role in scaffolding centrin filaments, we also investigated Sfi1 in cellular contraction and observed that some Sfi1 genes play a more prominent role than others. In particular, knockdowns of specific Sfi1 genes led to impaired contractile responses, including either complete failure to contract in response to a stimulus, or a marked reduction in contraction speed. These findings suggest that Sfi1 proteins in this system may have specialized roles in coordinating structural patterning, scaffolding oral primordium growth, and calcium-mediated contractility.

## Results

### Regional differences in centrin patterning on the Stentor cortex

*Stentor,* like other ciliates, has over 100 homologs of centrin and centrin-like proteins. Western blots using the centrin antibody 20H5 show that *Stentor* has centrin and isoforms at the average typical size (20-25kDa) as well as predicted large calmodulins (55-70kDa) and other EF-hand containing proteins of varying molecular weights (Figure 1B). To characterize the plethora of centrin homologs, we first used maximum likelihood to infer a phylogenetic tree with characterized human, yeast, and *Paramecium* centrin and calmodulin sequences as hallmarks. 36 of 80 potential centrin candidates in the *Stentor* genome that were also found in the set of EF-hand proteins identified by proteomic analysis of *Stentor*^39^, aligned well to the hallmark sequences common to centrins in other organisms. On our tree, only one sequence (SteCoe_3383) falls into the group of canonical yeast and human centrin3-like proteins that are found in most eukaryotes (Figure 1C). 18 sequences group together with *Paramecium* centrins thought to be ciliate-specific, like the ones found in the infraciliary lattice^40^. 16 sequences fall in the highly diverged group of calmodulin-likes and may not represent bona fide centrin proteins. One sequence (SteCoe_24203) sits in a group sister to centrin3-likes.

To visualize the centrin cytoskeleton, we performed immunofluorescence of centrin using a commercial monoclonal antibody, coupled with expansion microscopy. An expansion factor of approximately 9.2 was calculated from measurements of unexpanded cells and post expanded cells taken from the same culture. Our results show localization in the cortical fibers that run parallel along the cell, particularly in the posterior, showing dense localization with tapering intensity moving towards the anterior of the cell (Figure 1D). In addition to localization in the cortical fibers, the anterior of the cell contains a mesh-like network of centrin with a striking boundary halfway through the cell towards the posterior. When looking at a single z-slice of centrin immunofluorescence using conventional confocal microscopy, ladder-like rungs can be seen in the posterior of the cell, which do not appear in the maximum intensity projections, which is due to the much greater intensity of the myoneme bundles (Figure 1E). This complex pattern raises the question of what is organizing centrin and what makes centrin different in the anterior and posterior. Centrin binding proteins have been shown to have roles in modulating the activity of centrin in different processes such as ciliary and flagellar function and cell cycle regulation^41^. Sfi1, with its many centrin binding repeats and involvement in the myonemal cytoskeleton in other ciliates^34,35,42^, is a strong candidate for organizing centrin.

### Stentor contains multiple Sfi1 encoding genes

As with the centrin family, the Sfi1 gene family is also expanded in *Stentor*, with at least 15 members based on motif detection using RADAR and ScanProsite (Table 1)^43,44^. The largest Sfi1s have high degrees of homology with up to 50 potential repeats, found through searching for repeated motifs ending with W[KR], which is much greater compared to the 23 repeats found in the human homolog^28^, and consistent with ciliate Sfi1s have many more repeats ^30,32,35^ compared to humans and yeast. A consensus sequence for the Sfi1 motif was generated in *Stentor* using repeats found in SteCoe_17904, SteCoe_25973, and SteCoe_10021. This consensus sequence in *Stentor* is more similar to those found in other ciliates, in particular *Spirostomum minus*, compared to humans and yeast, consistent with the predicted phylogeny. While the sequence still contains the conserved tryptophan at the 22^nd^ position and R/K at the 23^rd^ position (Figure 2B), the initial residues appear to be less conserved, with many repeats beginning with a KGAL sequence. The previously reported presence of proline and glycine resides in the sequence signature found in *Spirostomum*^35^, which are predicted to create kinks in the alpha-helical structure, are also found in the sequence signature of *Stentor* Sfi1.

**Table 1:**

| Accession | MW<br>(kDa) | RNAseq<br>cluster | MS |
| --- | --- | --- | --- |
| SteCoe_17904<br>(Sfi1-1) | 434.3 | 2 | A |
| SteCoe_25973<br>(Sfi1-2) | 527.27 | 4 | A |
| SteCoe_10021<br>(Sfi1-3) | 495.27 | 5 | A |
| SteCoe_8740 | 239.13 |  | P |
| SteCoe_17381 | 241.25 | 2 |  |
| SteCoe_3113 | 309.55 |  | A |
| SteCoe_24260 | 140.78 |  |  |
| SteCoe_20357 | 354.87 |  |  |
| SteCoe_10583 | 360.9 |  |  |
| SteCoe_8584 | 184.41 | 4 |  |
| SteCoe_20206 | 183.51 | 5 | A |
| SteCoe_7525 | 152.92 | 3 |  |
| SteCoe_30007 | 179.45 |  | P |
| SteCoe_3004 | 112.07 |  |  |
| SteCoe_10143 | 143.2 |  |  |

**Figure 2:**
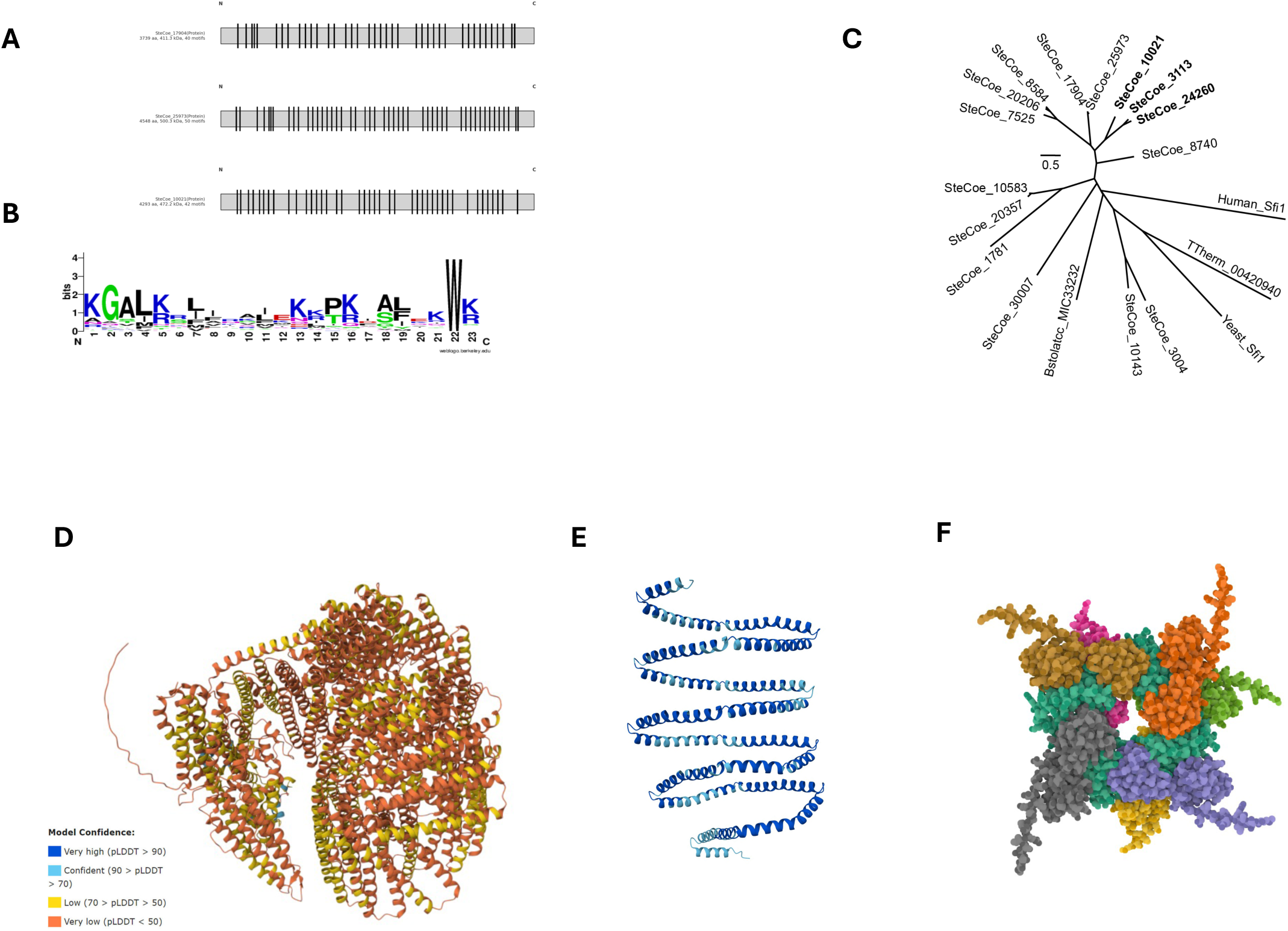
**Multiple Sfi1 encoding genes are found in *Stentor*** (A) Schematic of the three largest Sfi1s in *Stentor:* SteCoe_17904, SteCoe_25973, and SteCoe_10021, and their putative centrin binding motifs found from RADAR repeated motif searches ending in W[KR] (B) Sequence logos of Sfi1 repeats in *Stentor* generated using Sfi1 repeats from SteCoe_17904, SteCoe_25973, and SteCoe_10021 using the tryptophan at the 22nd position as a diagnostic residue. *Stentor* sequence mostly closely resembles that from *S. minus*, containing helix-breaking residues alanine and glycine and beginning with KGAL in positions 1-4^34,35^. (C) Maximum likelihood phylogenetic tree of Stentor Sfi1 homologs showing their relationship to Sfi1 homologs in other organisms including Human, Tetrahymena (TTherm), Yeast and Blepharisma (Bstolatcc). Branch-lengths represent average amino acid substitutions per site. (D) Alphafold3 prediction of full length SteCoe_17904 shows low model confidence. (E) Alphafold3 prediction of 7 Sfi1 repeats from SteCoe_17904. Sfi1 repeats are largely alpha-helical, and kink at evenly spaced repeats, creating a coil. (F) Alphafold3 prediction of 7 Sfi1 repeats from (E) bound to 7 centrin3. The centrins, each a different color, bind to the coiled Sfi1 backbone (teal). Diagnostic tryptophan residues are highlighted (green, right panel), indicating the site of centrin binding.

Analysis of the Sfi1 sequences revealed a series of internal repeats with the consensus motif separated by gaps of 46 amino acids. This gap is larger than that seen in Sfi1 motifs in humans and yeast. While *S. cerevisiae* Sfi1p repeats vary in length between 23-36 amino acids^29^, *Stentor* Sfi1 repeats are more consistently evenly spaced apart, as shown by the centring binding W[KR] site distribution (Figure 2A). Because of the large size of the entire Sfi1 proteins and the disordered tails at the N and C terminals, we are unable to predict structures of entire Sfi1 proteins with high confidence (Figure 2D). *Stentor* Sfi1 protein structure is predicted to be mainly alpha-helical, as confirmed by an Alphafold3^45^ prediction of seven Sfi1 repeats from SteCoe_17904 (Figure 2E). As the yeast Cdc31p centrin 3 is known to bind to yeast Sfi1, we also used Alphafold3 to predict the structure of these seven Sfi1 repeats bound to seven molecules of *Stentor* centrin 3, which creates a super-helix forming filaments or fibers (Figure 2F). Previously, sequence and structural analyses in *S. cerevisiae* show individual centrins stabilized by head-to-tail contacts form a helical filament surrounding the central Sfi1 alpha helix, with spacing between adjacent repeats that varies between 23-33 residues^29^. In contrast, in *Stentor* and *Spirostomum*^35^, the organization of centrin-binding regions are highly regular over the majority of the protein’s length, containing consecutive 69-amino acid repeats (23 amino acids of the Sfi1 motif plus 46 amino acids before the next motif repeat). In *Stentor*, the centrins are predicted to bind to the core Sfi1 alpha helix, creating a supercoil of alpha-helices, where each turn of the superhelix contains 4 centrin binding sites.

With a large number of Sfi1 family members, we applied two criteria to identify family members most likely to have functions related to spatial patterning. First, we leveraged a strength of *Stentor* in its ability to regenerate patterning after perturbation. To do so, we turned to a transcriptomic analysis of regeneration to identify the candidates most likely to play a role in this process. Multiple Sfi1s are upregulated in various timepoints during the transcriptional program of *Stentor* oral regeneration (Table 1)^46^. As a second criterion, we leveraged a previous proteomic study to look for Sfi1 family members with significant localization to the anterior of the cell versus the posterior, some Sfi1s are found to be more abundant in the anterior vs the posterior and vice versa^39^, indicating that some, but not all, are more localized to a specific half of the cell and may perform different functions within the cell (Table 1).

We found that the three largest Sfi1 proteins in the *Stentor* genome, SteCoe_17904, SteCoe_25973 and SteCoe_10021, were all upregulated during oral regeneration, and also showed regionalized enrichment along the anterior-posterior axis, and we therefore selected these three for further analysis. From here on, SteCoe_17904, SteCoe_25973 and SteCoe_10021 will be referred to as Sfi1-1, Sfi1-2, Sfi1-3 respectively.

To determine the localization of Sfi1 in *Stentor*, we generated separate polyclonal antibodies against each of the three proteins Sfi1-1, Sfi1-2, and Sfi1-3. Because of the high level of homology among the repeats in the three proteins, peptides taken from the N-terminal region, outside the region of the repeated motifs, were used to generate the antibodies to each of the three proteins. In Sfi1-2 and Sfi1-3 immunoblots, we see that the antibodies detected large proteins in the expected size range (527.3kDa and 495.7kDa for Sfi1-2 and Sfi1-3 respectively). The Sfi1-1 antibody (Figure 3A) detected a distinct band at a high molecular weight consistent with the predicted molecular weight of the Sfi1-1 protein (434.3kDa, Figure 3B, arrows) but also detected a number of significant non-specific protein bands. Additionally, there is staining detected in the aggregated material remaining at the bottom of the loading well (Figure 3B, asterisk). Because of the large size of these proteins, they may not all be able to successfully migrate through the gel. Non-specific bands may be due to cross-reactivity because of the high level of homology among Sfi1 proteins or partial degradation of protein due to proteases that may still be present in the cell lysate. Separate blots for each antibody are shown in Supplementary Figure 1. To confirm specificity of the antibodies in immunofluorescence experiments, we used a peptide block with corresponding peptides used for antibody production. Antibodies were incubated with their corresponded peptides and immunofluorescence was performed using these pre-incubated antibodies, which led to a virtually complete loss of Sfi1fluorescence intensity per cell for all three antibodies (Supplementary Figure 2).

**Figure 3:**
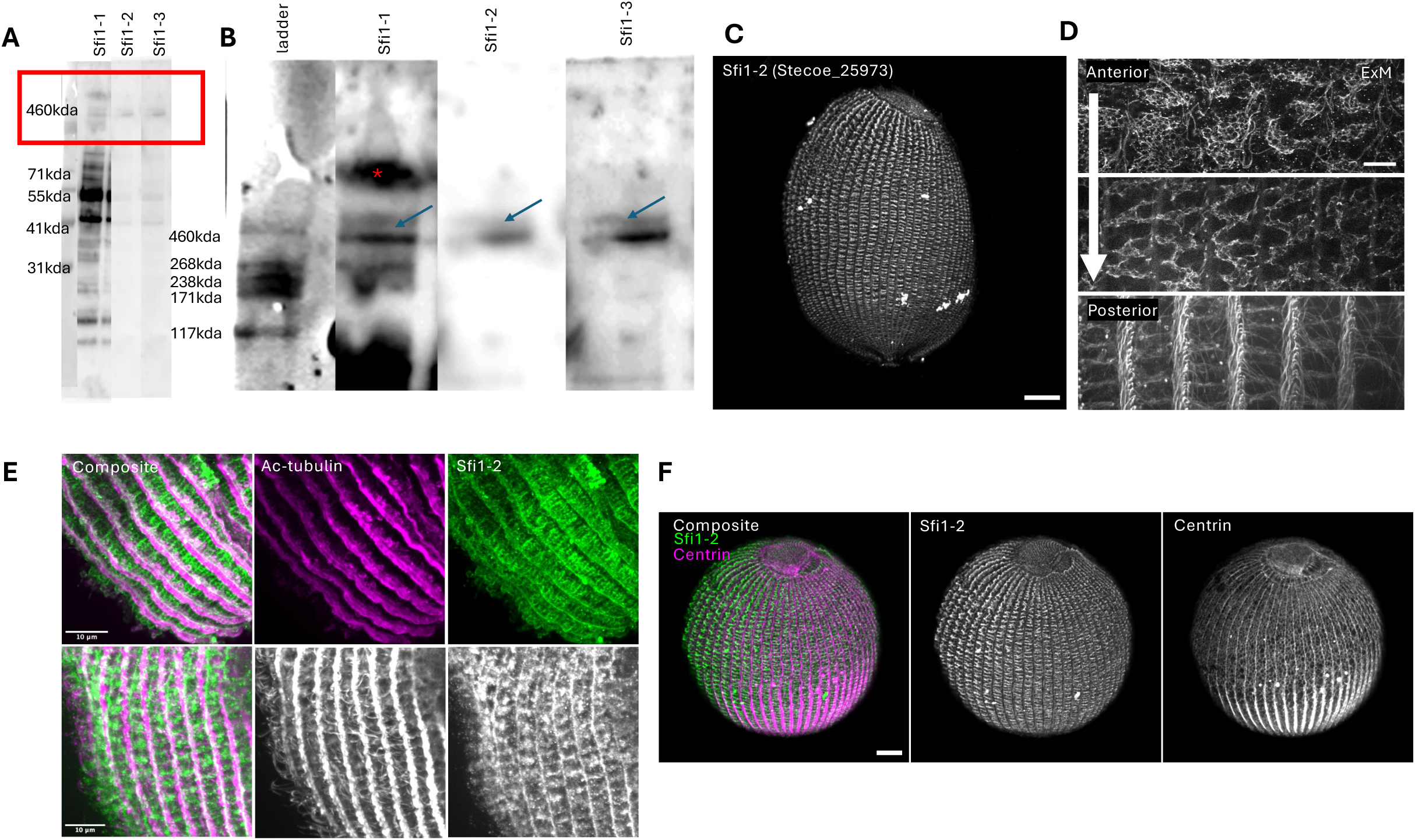
Sfi1 antibody localizes to distinct structures. (A) Western blot of custom Sfi1-1, Sfi1-2, and Sfi1-3 antibodies on *Stentor* lysates. Faint protein bands are visible at a high molecular weight corresponding to estimated sizes of Sfi1 proteins. Sfi1-1 contains many non-specific bands, which may be due to protease activity during lysate preparation, the polyclonal nature of the antibodies, and high homology of Sfi1 proteins. (B) Western blot close-up of high MW area, red asterisk indicates lysate aggregate at the bottom of the well. Large proteins are detected in the estimated size range of *Stentor* Sfi1 (blue arrows) (C) Representative image of maximum intensity projected confocal z-stacks of a cell stained with Sfi1-2 antibody. Anterior irregular bands gradually become more distinct and regular moving posteriorly along the cell. (D) Representative max intensity projection images of confocal z-stacks on expansion microscopy treated cells showing Sfi1 anterior mesh (top) and mid-anterior bands (middle) shown with Sfi1-2 immunofluorescence. Moving posteriorly, the bands become more organized and regular (bottom) as shown with the general protein stain NHS ester. Scale bar 1um. (E) Representative max intensity projection images of confocal z-stacks of Sfi1-2 (green) co-labeling with acetylated tubulin to reveal cilia (magenta). The bands of Sfi1 in between the cortical rows do not co-localize with cilia but rather lie beneath the cilia. Scale bar 10um. (F) Representative images of max intensity projection of confocal z-stacks of centrin (magenta) and Sfi1-2 (green). The anterior centrin mesh does not co-localize with the Sfi1 bands, however there is some colocation given that the Pearson’s correlation coefficient equals 0.42, n=36. Scale bar 10um.

Immunofluorescence was also eliminated by RNAi knockdown of each of the Sfi1 genes as discussed below (Supplementary Figure 5). Immunofluorescence of all three Sfi1 antibodies showed a similar pattern (Supplementary Figure 3), with wide disorganized filaments at the anterior of the cell and more organized, thin, ladder-like rungs in the posterior, as well as tapering cables from the posterior to the anterior (Figure 3C). To take a closer look at the differences between the Sfi1 organization in the anterior and posterior of the cell, we turn to expansion microscopy to expand our samples by up to 10-fold. Because Sfi1 custom antibodies are not optimized for expansion microscopy, they produced a very weak fluorescent signal especially in the posterior of expanded cells (Supplementary Figure 4, bottom middle panel). In double-labeling experiments we found that several structures stained with Sfi1 antibodies also stained with the general protein stain NHS-ester (Supplementary Figure 4). Specifically, the parallel cables that run along the cell, the ladder-like rungs perpendicular to the cables in the posterior, and a banded mesh network in the anterior. The fact that these structures show up with NHS-ester, which labels all protein, suggests that Sfi1 proteins are found within protein-dense structures. We note in particular that the posterior ladder like rungs seen by NHS-ester staining all stain with Sfi1 antibodies. Moreover, within the filamentous network composing these lateral elements, there is perfect co-localization of Sfi1 with NHS fluorescence. Sfi1 antibodies also stain punctate structures outside of the lateral filaments that do not correlate with NHS staining. The high correlation of Sfi1 immunofluorescence and NHS staining within the lateral filaments means that we can image NHS-ester fluorescence to visualize the detailed structure of the Sfi1-containing posterior lateral filaments. A 10x expansion method^47^ reveals bands of a mesh network within the anterior bands (Figure 3D, top panel), distinct from the centrin mesh-like network in that the mesh is contained within the lateral bands. Each filamentous element composing the mesh measures approximately between 0.73µm-2.4µm corresponding to 1-4 Sfi1 proteins based on the estimated length of 580nm for an Sfi1 superhelix (Figure 2E). Moving posteriorly, the mesh-like filaments become more organized (Figure 3D, middle panel), gradually forming a ladder rung organization in the posterior (Figure 3D, bottom panel).

The ladder-rung like posterior bands appear to correspond to the connections between M-bands shown in early illustrations of *Stentor* fiber systems^6,10^ in which perpendicular connections connect adjacent longitudinal myonemes. The thickness of the posterior bands spans a length corresponding to several basal bodies in the kinety, such that for every band, there are approximately 2-5 basal body pairs corresponding to 1.7-4µm. The lateral bands appear to initiate at the basal bodies, and their length spans the width of the gaps between adjacent myonemes, which corresponds to the pigment stripes (Figure 3D) ranging from 5-13µm depending on the circumferential position of the stripes. The spacing between the bands is not consistent but ranges between 3-9µm. The shortest rungs are also the thinnest and increase in thickness with length up to a certain point, after which the thickness can vary. We note that these lateral bands are distinct from cilia as shown by double-labelling with an acetylated tubulin antibody (Figure 3E). They lie beneath the cilia, consistent with the z-position of the “M-band connectives” described by Tartar^6^. In contrast with the lack of co-localization of the Sfi1 proteins with centrin in the anterior centrin mesh, in the posterior lateral bands Sfi1co-localizes with centrin (Figure 3F, Figure 1E) with a Pearson’s correlation coefficient of 0.42 determined using the plugin JACoP (https://imagej.net/plugins/jacop) in ImageJ, n=36. Because these Sfi1 co-localize with centrin specifically in the posterior region of the cell, this suggests a region-specific interaction that may reflect functional specialization of the cytoskeletal scaffold along the longitudinal axis, although the possibility that a different Sfi1 co-localizes with the anterior centrin mesh cannot be ruled out.

We conclude that the three Sfi1 proteins selected for this study co-localize with the myonemes and with lateral centrin bands in the posterior part of the cell, but not with the anterior centrin mesh, suggesting they may play a specific role in forming a sub-set of centrin structures.

### Role of SFI1 family proteins in maintaining anterior/posterior cortical patterning of centrin

We next tested the functional role of the three Sfi1 family members in cortical patterning of centrin. Cells were microinjected with in vitro transcribed dsRNA directed against each of the Sfi1 family members. Regions of 1.3-1.5kb of each Sfi1 were targeted by RNAi using sequences specific to each Sfi1(Supplementary Table 1). In all three cases, protein depletion was verified by immunofluorescence with the corresponding Sfi1 antibodies. In the case of RNAi mediated knockdown of Sfi1-1 and Sfi1-2, cells stained with anti-centrin antibodies revealed the loss of the distinct anterior and posterior differences in centrin patterning. In both cases, the mesh-like network normally seen only in the anterior of control cells now appears in both the anterior and the posterior, while the thick myonemes normally seen in the posterior show notably reduced centrin incorporation (Figure 4A). In Sfi1-3 RNAi cells, centrin retains its distinct anterior/posterior patterning. We quantified this observation by measuring the textures of the anterior and posterior across 3 points in each cell using the GLCM texture analyzer in ImageJ and took the ratio of the posterior over the anterior, where a greater ratio indicates more distinct textures (Figure 4B). In addition to the loss of posterior structures in the Sfi1-1 and Sfi1-2 RNAi cells, we also found that the mesh-like network in the anterior is much less dense and appears disassembled or disrupted compared to control cells (Figure 4C). Thus, both the anterior and posterior halves of the cell show abnormalities in centrin distribution, with the main phenotype being a reduction in the regional differences in patterning between the two halves. The overall integrated fluorescence intensity of centrin immunofluorescence staining is not significantly different in the RNAi cells compared to controls, showing that the distribution is altered but not its overall abundance (Figure 4D). The fact that the overall centrin fluorescence is not affected in RNAi knockdowns is consistent with distinct Sfi1 isoforms playing site-specific roles in patterning rather than globally affecting centrin gene expression or protein stability.

**Figure 4:**
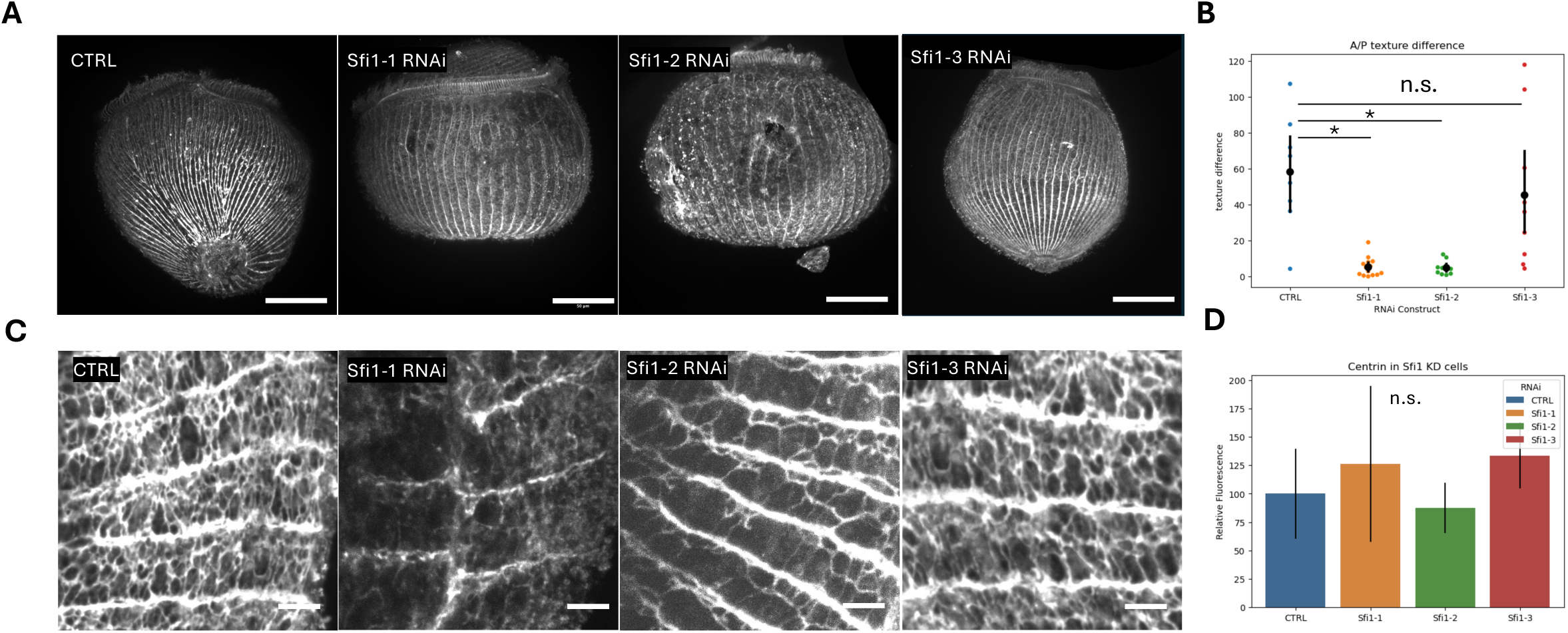
Role of Sfi1 family proteins in maintaining anterior/posterior cortical patterning of centrin. (A) Representative images of max intensity projection of confocal z-stacks of centrin in control and Sfi1 RNAi knockdowns. The anterior mesh network of centrin in Sfi1-1 and Sfi1-2 appear disrupted compared to control and Sfi1-3. Scale bar 50um. (B) Plot showing the texture difference in centrin immunofluorescence images from Sfi1 knockdown cells. Three sections of the anterior and three sections of the posterior of each cell was analyzed using a GLCM texture analysis. The average ratio of the posterior texture over the anterior texture was calculated for reach condition. A lower texture difference indicates that the anterior and posterior textures are more similar. Bars indicate the mean texture differences and lines indicate S.E.M. Control 58.3±31.7, n=9, 5.3±5.26 for Sfi1-1 n=14, 4.9±3.7 for Sfi1-2 n =11, and 45.4±39.4 for Sfi1-3 n=11. Statistical significance was calculated using a student’s t-test where *p<0.05 (C) Representative images of max intensity projection of confocal z-stacks of anterior centrin mesh in control and Sfi1 RNAi knockdowns. The mesh in Sfi1-1 and Sfi1-2 knockdowns is much less dense with less branching tham that of controls and Sfi1-3. Scale bar 1um. (D) Plot showing the corrected total cell fluorescence of control and Sfi1 knockdown cells stained with centrin. Fluorescence intensities of RNAi cells are normalized to that of the control and bars indicate mean whole cell fluorescence and lines indicate S.E.M. Average fluorescence for Sfi1-1 RNAi knockdown is 126.2±68.5, for Sfi1-2 87.68±22.2, and for Sfi1-3 133.7±28.8. The total fluorescence intensity is not significantly affected in the Sfi1 knockdowns (p>0.05, n=10 for each condition) compared to the control cells. Statistical significance calculated with a student’s t-test.

### Sfi1 proteins required for progression of oral regeneration

Classically, *Stentor* has been used as a model for regeneration, due to its amenability to microsurgical manipulations, and robust regeneration assays^6^. When exposed to a sucrose shock or bisected with a glass needle, *Stentor* will regenerate a new oral apparatus over the course of 8-10 hours at room temperature. Oral regeneration begins with assembly of basal bodies and associated structures at the oral primordium site, where the narrow and wide cortical cables intersect. This precise location on the ventral surface of the cell is at a considerable distance from the position of the original oral apparatus. The basal bodies eventually form an oral primordium which elongates along the cell, curls upward at the posterior end to form a new buccal cavity or gullet, and finally migrates to the anterior of the cell to the normal position occupied by the membranellar band (Figure 5A). While previous studies have produced a transcriptional profile of regeneration and a proteomic analysis^39,46,48,49^, much less is known about specific molecular players controlling this complex morphological process. Microsurgical experiments have shown that the signal for the initiation of regeneration travels over the cortex^50^. Local anesthetic such as tetracaine and dibucaine which act on the cell membrane and affect calcium metabolism have been shown to delay oral regeneration, specifically in early stages when much of the primordium is being built^51^. This delay might then be caused by the disruption of cytoskeletal elements involved in transmitting the stimulus for oral primordium formation. Given that oral regeneration involves formation and alignment of thousands of basal bodies and requires a signal that travels through the cytoskeleton and that might involve calcium binding proteins, we reasoned that centrin and Sfi1 proteins might play a role in regeneration.

**Figure 5:**
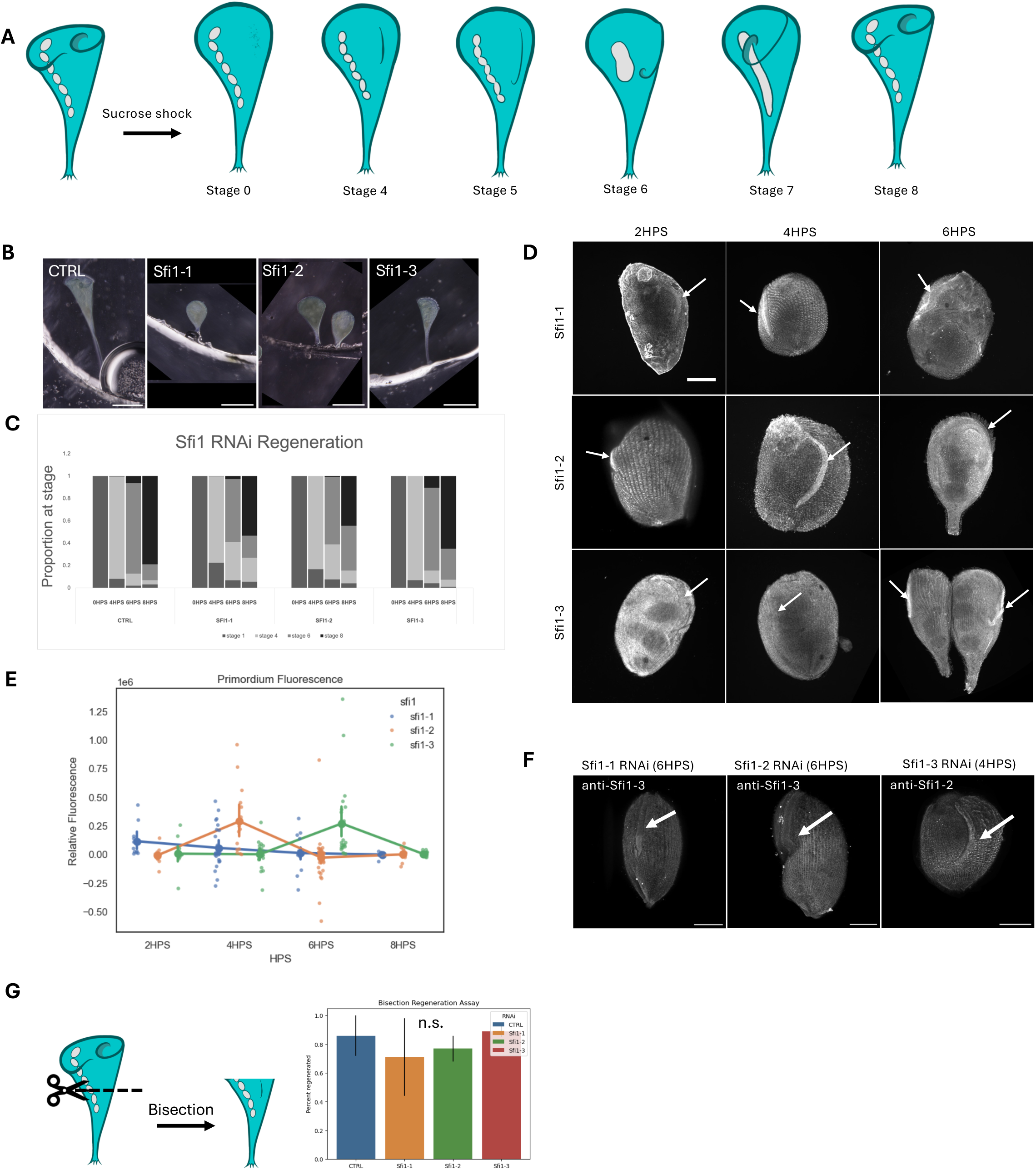
Sfi1 is required for regeneration after sucrose shock. (A) Regeneration assay in *Stentor*. Each stage corresponds to hours post sucrose shock. Sucrose shock is used to induce shedding of the membranellar band. Once removed, the cell regenerates a new oral apparatus, first with basal bodies assembling at the primordium site. The primordium elongates along the length of the cell (Stage 4) and curves at the posterior end to form the gullet (Stage 6). This structure then migrates to the anterior of the cell to the position of the original oral apparatus (Stage 8). (B) Representative brightfield images of *Stentor* cells 8 hours post sucrose shock. Membranellar bands are visible at the anterior of the control and Sfi1-3 RNAi cell, having a conical shape. Sfi1-1 and Sfi1-2 RNAi cells are missing the ring of cilia at the anterior, creating a rounded, tear-drop shape. Scale bar 100um. (C) Plot showing the percent of cells that have reached the corresponding stage of regeneration at the specified timepoints. A larger proportion of Sfi1-1 and Sfi1-2 RNAi cells remain in stages 4-6 compared to control and Sfi1-3 RNAi cells. Data was generated from 5 independent trials, each with n=20-120. (D) Representative images of maximum intensity projection of confocal z-stacks showing a time course of oral primordium growth during regeneration with immunofluorescence of corresponding Sfi1 antibodies. Arrows indicate the oral primordium. Scale bar 50um. (E) Plot depicting the fluorescence intensity of Sfi1 in the primordium at different timepoints. Sfi1-1 intensity remains relatively constant throughout regeneration, however, Sfi1-2 peaks at 4HPS, and fluorescent is largely absent from the oral primordium at 6HPS, as seen in (D). In contrast, Sfi1-3 has fluorescence visible in the primordium only at 6HPS, although the fluorescence of the whole cell across timepoints remains relatively constant. Bars indicate mean fluorescence intensity. Sfi1-1 n=14, 17, 27, 18; Sfi1-2 n=33, 25, 35, 20; Sfi1-3 n=20, 16, 38, 31 for each respective timepoint. (F) Representative images of maximum intensity projection of confocal z-stacks showing little Sfi1-3 fluorescence in the oral primordium of Sfi1-1 and Sfi1-2 knockdown cells (left, middle panels). Right panel shows Sfi-2 fluoreescence in the primordium of Sfi-3 RNAi knockdown cells. (G) Plot showing percent of Sfi1 RNAi cells and controls that completed regeneration in 8 hours post bisection with a glass needle. Bars indicate mean proportion of cells that regenerated, and lines indicate S.E.M. with control 0.86±0.14, Sfi1-1 0.71±0.27, Sfi1-2 0.77±0.09 and Sfi1-3 0.89±0.02. Three independent trials were performed, each with n=20-35 cells. The percent regenerated across all conditions is not statistically significant, as calculated using a student’s t-test with p>0.05.

The three Sfi1 proteins analyzed in this study were all encoded by genes showing upregulation during oral regeneration^46^. However, their timing of expression was different. Our previous transcriptomic analysis indicated five distinct clusters of gene expression based on the timing of peak expression. The earliest expressed cluster was designated cluster 2, with clusters 3-5 peaking at progressively later times during regeneration. Sfi1-1 is upregulated early in regeneration, in cluster 2, which is also the time of peak expression of centriole biogenesis related genes and corresponds to the time when new basal bodies, that will ultimately assemble into the membranellar band, first start to appear. Sfi1-2 peaks several hours later in cluster 4, which corresponds to the time at which motile cilia have assembled onto the basal bodies and the cilia have begun to align into the structure of the membranellar band. Sfi1-3 peaks later in regeneration in cluster 5, the time when the oral primordium has fully assembled and the cell is now re-organizing its microtubule fibers anterior of the oral primordium to place it at the anterior end of the cell.

To test the role of Sfi1 family members in regeneration, each of the three Sfi1 genes were knocked down by RNAi and then the cells were induced to regenerate using a sucrose shock which removes the oral apparatus. RNAi was confirmed to result in reduction in protein quantity, using quantification of Sfi1 intensity in immunofluorescence images (Supplementary Figure 5). The Sfi1-1 and Sfi1-2 RNAi cells started to regenerate, forming a membranellar band, but did not fully regenerate and remained arrested in stages 4-6, such that the oral primordium formed but did not complete migration to the anterior of the cell (Figure 5B, Figure 5C). Additional timepoints at 24 hours post shock (data not shown) were taken to confirm that regeneration would not continue beyond the typical 8-hour time period. In Sfi1-3 RNAi cells, however, the cell is able to completely regenerate forming a fully normal looking membranellar band at the anterior of the cell. These findings suggest that the earlier expressed isoforms, Sfi1-1 and Sfi1-2, are required for the later stages of regeneration which includes the elongation and migration of the oral primordium along the cell.

Time course analysis of Sfi1 localization during regeneration revealed distinct temporal patterns for each isoform. Sfi1-1 staining in the oral primordium can be seen in early timepoints compared to 6HPS (Figure 5D), and while the fluorescence intensity remains relatively constant during regeneration, it does have slightly increased intensity early in regeneration (Figure 5E), consistent with Sfi1-1 gene expression early in regeneration^46^. Once the oral primordium becomes visible, usually at 4HPS, Sfi1-1 is seen in the primordium (Figure 5D, arrow) as shown by the increase in immunofluorescence. However, by 6HPS much of that intensity is lost. In the case of Sfi1-2, fluorescence in the oral primordium peaks at 4HPS and with clear localization in the oral primordium and similarly shows reduced signal at 6HPS. In contrast, Sfi1-3 shows maximum recruitment to the primordium at 6HPS but not 4HPS (Figure 5D, Figure 5E). These data suggests that these Sfi1 proteins are sequentially recruited to the primordium, with timing consistent with their transcript expression during regeneration. This sequential localization pattern supports the idea that Sfi1 isoforms are differentially and temporally regulated. We next asked if earlier scaffolding Sfi1 are required for the recruitment of later expressed Sfi1. When we look at Sfi1-3 immunofluorescence in Sfi1-1 and Sfi1-2 RNAi knockdown cells, we see an absence of signal in the oral primordium (Figure 5F). However, if we look at Sfi1-2 intensity in Sfi1-3 RNAi knockdown, we do see signal in the oral primordium (Figure 5F, right panel). These data suggest that not only are different Sfi1 isoforms sequentially localized to the oral primordium, but their recruitment requires prior recruitment of earlier Sfi1 isoforms to the regenerating oral primordium, indicating a sequential scaffolding mechanism.

These time course images also revealed a loss and re-establishment of Sfi1 bands during regeneration. In the first 3 hours of regeneration (Supplementary Figure 6), the lateral bands become more punctate until eventually fluorescence occurs only in the longitudinal cables. The punctate bands re-appear and eventually resume their normal appearance near the end of regeneration. We speculate that this changing pattern of Sfi1 protein localization could be due to a temporary shift of Sfi1 from the ladder rung-like bands of the cytoskeleton to the oral primordium during regeneration.

The foregoing analysis was based on inducing oral regeneration by sucrose shock, which causes the cell to shed its existing oral apparatus including the membranellar band, although it retains the frontal field (region of the anterior bounded by the membranellar band). When Stentor cells are bisected with a glass needle, the anterior half-cell retains the membranellar band, but the posterior half-cell has to regenerate a new one. Interestingly, Sfi1 RNAi cells triggered to regenerate using bisection with a needle, rather than with sucrose shock were fully able to complete all stages of regeneration of the oral apparatus (Figure 5F). This is consistent with the fact that Sfi1 transcripts are upregulated in the sucrose shock response to regeneration but not the bisection response^46^. Potential differences between the two regeneration paradigms include the fact that the frontal field is removed in bisection but not in sucrose shock, or that the bisected cell will make a much smaller oral apparatus, proportional to its new body size^7^, reducing the need for basal body biogenesis. In any case, the fact that RNAi of Sfi1 family members expressed during regeneration in sucrose shock but not bisected cells, has an effect on the process in the former but not the latter case, further supports the specificity of the RNAi phenotypes observed.

### Role of Sfi1 in cellular contractile machinery

We have shown that Sfi1 isoforms play a role in patterning centrin on the cell cortex. This cortical centrin patterning is visually striking and structured, but does it have any physiological function? One possibility is that this centrin organization contributes to the mechanical properties of the cell, such as its contractility. *Stentor* is capable of contracting from an extended trumpet shape to a compact sphere on a scale of milliseconds (Figure 6A). This dramatic reduction in total body length is driven by specialized contractile fibers known as myonemes. These myonemes undergo a dramatic change in morphology of the filaments upon stimulation by intracellular calcium influx, coiling into helical structures^24,25^. The myonemes rapidly shorten in length and thicken in diameter. When relaxed, the myonemal filaments have a diameter of 40Å, but once contracted the diameter is increased to 120Å^37^. When contracted, the bundles of microfilaments that make up the myonemes are straight, but when the cell is extended, they become sinuous folds^36^. This transition enables both force generation and spatial reconfiguration of the cell body. Given that centrin has established roles in other ciliate contractile systems^30,34,35,42,52^, it is possible that the patterned organization of centrin could serve to modulate myoneme activity, ensuring that contraction occurs in a coordinated manner across the large surface of the cell.

**Figure 6:**
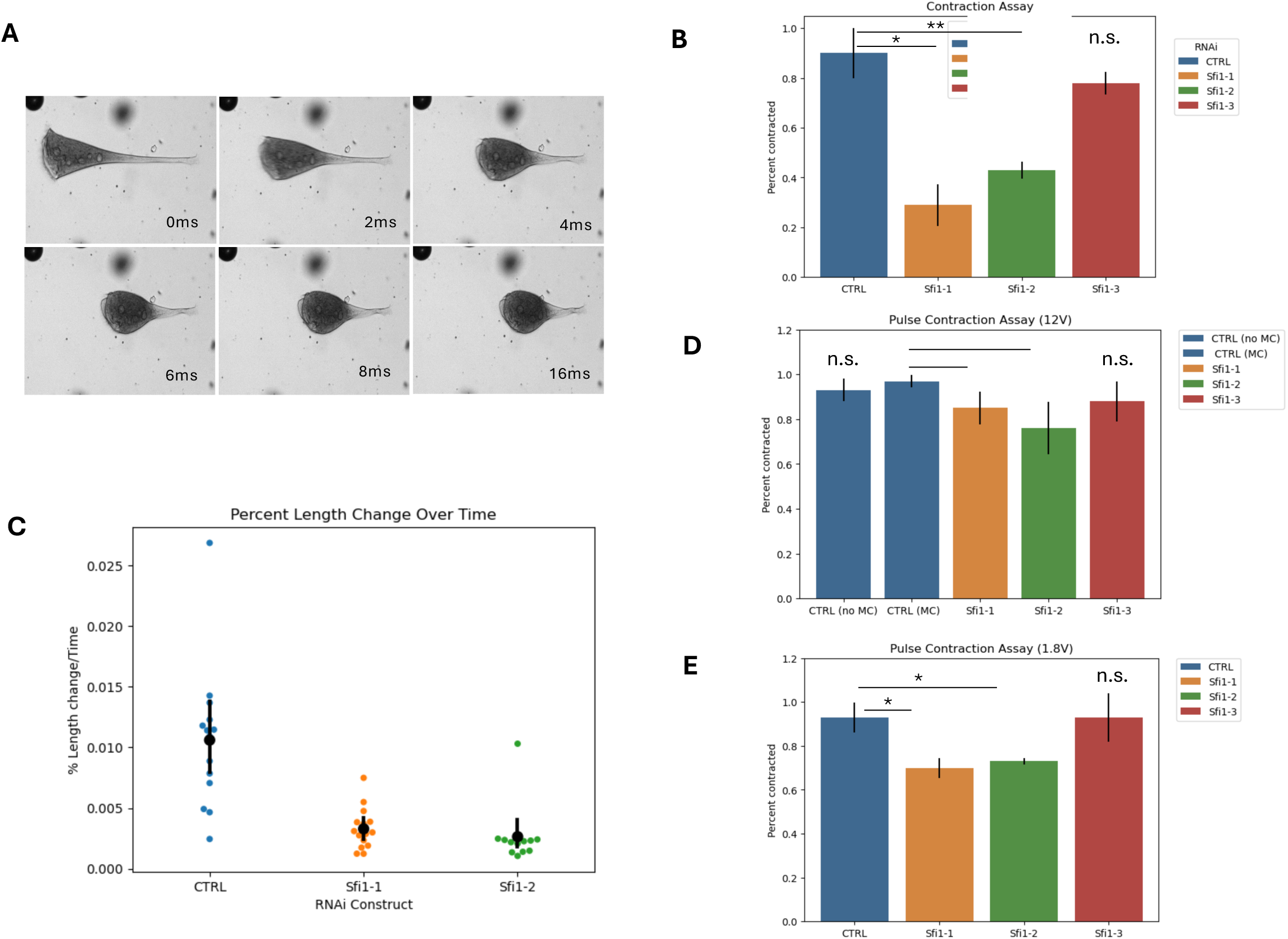
Role of Sfi1 in contractile machinery. (A) Representative still frame images of a contracting *Stentor* cell. Contraction occurs over timescale of milliseconds. (B) Plot showing the percent of cells that contract in response to the mechanical stimulus of being pipette up with a P20 pipette. Data was collected over 3 independent trials, with n=50 for each condition. *p<0.05, **p<0.005 (Sfi1-1, p=0.0015, Sfi1-2 p=0.00026, Sfi1-3 p=0.11). (C) Plot showing the percent of length change over time ratio for each condition. The average percent of length change over time for control cells is 0.82±0.59% per millisecond. For Sfi1-1, the average is 0.27±0.17% and for Sfi1-2, 0.20±0.04% per millisecond. (D) Plot showing the percent of cells that contract in response to a constant pulse of 12V. No statistical significance between control cells in spring water (CTRL no MC) and controls in 1.5% methylcellulose (CTRL mc, p=0.26). Sfi1-1 and Sfi1-2 show a decrease in contractility as compared to controls in methylcellulose. *p<0.005 (Sfi1-1 p=0.005, Sfi1-2 p=0.006, Sfi1-3 p=.068). (E) Plot showing the percent of cells that contract in response to a constant pulse of 1.8V. Sfi1-1 and Sfi1-2 show decreased contractility compared to control cells. *p<0.05 (Sfi1-1 p=0.015, Sfi1-2 p=0.025, Sfi1-3 p=0.48)

To test the role of Sfi1 in contraction, we knocked down Sfi1 using RNAi and stimulated contraction by pipetting extended cells into a P20 pipette tip. In four trials each with n∼50, 86.7%±4.5 of control cells contracted in response to the pipetting stimuli. Sfi1-1 knockdown cells, (24.3%±8.4) and Sfi1-2 knockdown cells (46.3%±3.4) responded to the stimulus much less than the controls. 72%±10.4 of Sfi1-3 knockdown cells contracted, showing a less severe contraction phenotype. These results indicated that Sfi1 RNAi cells are still able to contract, but did not indicate whether they contract at the same speed as controls.

Because *Stentor* contract on a scale of milliseconds, we turn to high-speed imaging at 5000 frames per second. Cells were gently flattened and allowed to extend on a 9mm spacer on a glass slide and contraction was stimulated with a mechanical force in the form of a gentle tap on the slide. Percent changes in cell length were not statistically significant between controls (n=14, 45.1±7.6) and Sfi1-1 (n=15, 41.1±7.8), however, some Sfi1-2 knockdowns (n=13, 35.9±2.9) do not contract as dramatically. Rather than contracting into almost perfect spheres, they contract to a longer, teardrop shape (Supplementary Video 1). Upon further analysis, the most dramatic effect was on the amount of time between contraction initiation and termination, which increased in both Sfi1-1 and Sfi1-2 knockdowns. On average, control cells completed contraction in 55.5±35.6 milliseconds, whereas Sfi1-1 and Sfi1-2 knockdowns completed contraction in 152.44±69.9 and 180.1±72.65 milliseconds respectively. Thus, contraction speed is much less in these knockdowns compared to control cells (Figure 6C).

To confirm that these results are not dependent on inconsistencies in pipetting or mechanical stimuli, we repeated the analysis using electrical pulses, which can also stimulate contraction in *Stentor*^37^. *Stentor* cells were placed in 1.5% methylcellulose and in a spacer made from flat pieces of steel functioning as electrodes with a gap of approximately 18-20mm. A coverslip was placed on top and using a short pulse of 12V, cells were stimulated to contract. Images pre and post stimulus were taken and percent of cells contracted calculated for each experiment. Cellular contraction was not significantly affected in methylcellulose (n=30-39 over 4 trials, percent of cells contracted 96.8±2.7) as compared to contraction in spring water (n=16-31, 4 trials, 93.0±5.3. While the results are not as dramatic as the previously described assays, Sfi1-1 (n=20-50, 6 trials, 84.7±7.3) and Sfi1-2 (n=20-28, 6 trials, 76.5±11.7) knockdowns show a statistically significant reduction in number of cells that do not immediately contract after an electrical stimulus. Consistent with previous assays, Sfi1-3 (n=23-43, 6 trials, 87.8±9.4) knockdowns contract at a rate within the range of control cells. We also repeated this experiment using a much lower pulse of 1.8V, and observed similar results, where Sfi1-1 (n=16-29, 4 trials, 0.70±0.05) and Sfi1-2 (n=10-21, 4 trials, 0.072±0.14) knockdown cells had decreased contractility, while Sfi1-3 (n=12-34, 4 trials, 0.93±0.01 contracted comparably to control cells (n=15-29, 3 trials, 0.93±0.07). These results indicate that proper Sfi1 function is required for the rapid contraction mechanism in *Stentor*.

## Discussion

### Regionalized patterning by regionalized distribution of scaffolding proteins

Regionalized differences in protein localization may occur through various processes such as membrane trafficking, cytoskeletal organization, localized translation, signaling pathways, and post-translational modifications. These processes are coordinated to work together to establish and maintain localized expression of proteins. For instance, in neurons, AnkyrinG is essential for organizing ion channels in the axon initial segment, the compartment that initiates action potentials and maintains neuronal polarity^53^. The localized accumulation of AnkyrinG serves as a cue for the recruitment and assembly of complex molecular machinery in a site-specific manner. Without AnkyrinG, maintenance of the axon initial segment is lost, along with neuronal polarity^54^. In the case of Sfi1 and centrin in *Stentor*, one compelling possibility could be that Sfi1 acts as a spatially organized scaffolding protein that governs the subcellular localization and assembly of centrin complexes. Scaffolding proteins are known to serve as molecular organizers, bringing together specific binding partners in defined cellular regions to enable the formation of higher-order structures or signal transduction networks. In this context, Sfi1 may play a critical role in anchoring centrin molecules along the contractile cytoskeleton in a manner that is not uniform but rather regionally patterned across the anterior-posterior axis of the cell. By compartmentalizing centrin assembly through a polarized and temporally varying distribution of distinct Sfi1 scaffolds, the cell can exert spatiotemporal control over cytoskeletal assembly, enabling the development of functionally distinct cellular domains. Thus, the Sfi1 scaffold may not only provide structural support for centrin binding, but it may also serve as a developmental or functional organizer, encoding spatial information into the architecture of the contractile cytoskeleton. This interpretation is further supported by the observation that the disruption of Sfi1 leads to decreased centrin mesh density and integrity of centrin-based cytoskeletal structures.

The model in which regional differences in the spatial arrangement of centrin fibers are produced by regional differences in Sfi1 isoform composition, is appealing because it reduced a very difficult problem (how to generate different complex 3D arrangements of filaments) into a simpler problem (how to generate differences in abundance of different proteins). Essentially, this model invokes patterning at the level of a scalar variable (Sfi1 abundance) to explain patterning at the level of a higher dimensional function (organization of centrin into different types of filament-based structures).

We note that, in addition to its well-characterized centrin binding repeats, Sfi1 also contains numerous putative calmodulin (CaM) binding sites, suggesting a potentially broader regulatory role in the assembly or modulation of the cytoskeleton. Calmodulin is a highly conserved calcium sensor and signaling adaptor and can act as a protein linker and regulator of scaffold proteins^55^. It can also bridge between different domains on same protein or can physically link two identical or different target proteins together. Structurally, CaM is made up of two independently folded lobes with the capacity of binding two Ca^2^ ^+^ ions each. The two lobes are made up of a pair of EF-hands each and they are connected by a long and flexible linker region, allowing the molecule to adopt diverse conformations in response to calcium signaling. The presence of predicted CaM-binding sites in Sfi1 raises the possibility that Sfi1 may serve as a multi-valent scaffold, coordinating interactions not only with centrin but also with calmodulin in a calcium-dependent manner. This dual interaction potential could enable Sfi1 to integrate mechanical and calcium-signaling cues, modulating its structural or organizational role within the cytoskeleton. For instance, calmodulin might regulate the binding affinity or conformation of Sfi1-centrin complexes, or alternatively, act as a dynamic crosslinker between Sfi1 filaments or adjacent cytoskeletal structures. Supporting this hypothesis, previous studies using calmodulin antagonists, trifluoperazine and W-7, have been shown to delay oral regeneration in *Stentor*^56^, suggesting that calmodulin is involved in the control or formation of the oral apparatus. It could be possible that calmodulin is involved in scaffold remodeling or reorganization during regeneration. Preliminary immunofluorescence imaging using an anti-calmodulin antibody further reveals a subcellular distribution pattern similar to that of Sfi1, reinforcing the idea that these two proteins may colocalize or interact with each other. While these observations are suggestive, direct biochemical or structural evidence for a Sfi1-calmodulin interaction has not been determined. Given the similarities between CaM and centrin, it could also be possible that CaM antagonists can bind to centrin itself, as previously shown in *Euplotes*^57^. Additional experiments, such as co-immunoprecipitation, in vitro binding assays, or cryo-EM structural analysis, will be needed to determine whether calmodulin binds directly to Sfi1 or whether such binding induces calcium-dependent conformational changes in the Sfi1 scaffold. Establishing this interaction would provide a mechanistic link between calcium signaling and contractile cytoskeletal organization, further supporting the role of Sfi1 as an integrative scaffold in the functional polarization of *Stentor* cells.

### Comparison with other cell types

Prior work on Sfi1 has established its role as a conserved component of the contractile cytoskeleton across multiple ciliates. In *Paramecium*, Sfi1 has been identified as a structural component of the ciliary lattice, a cortical scaffold that supports the coordinated beating of cilia and likely contributes to maintaining cortical architecture^30^. Similarly, in *Spirostomum* it is involved in the contractile cytoskeleton, which drives its rapid and forceful contractions^34,35^. Additionally, in *Vorticella,* Sfi1 localizes to the spasmoneme, a contractile stalk that rapidly coils like a spring in response to stimuli^42^. These findings underscore a broadly conserved role for Sfi1 in contractile systems among ciliates, in coordination with centrin and related proteins. Here, we extend these observations by demonstrating that Sfi1 in *Stentor coeruleus* is also a component of the contractile cytoskeleton, but with a more complex and multifunctional role. In addition to supporting contractility, our data shows that Sfi1 is sequentially recruited to the oral primordium during regeneration, where it may function as a scaffold for oral primordium growth.

The centrin-Sfi1 complex appears to be conserved throughout many eukaryotes, and our findings support this for *Stentor*. The repeated motif characteristic of Sfi1 proteins contains the conserved tryptophan at the 22^nd^ residue, with each repeat binding to one centrin molecule. The predicted structure contains long alpha-helices arranged in a coiled-coiled structure. *Stentor,* like other ciliates, has more centrin binding repeats compared to humans and yeast. In yeast, Sfi1 localizes to the spindle pole body where it forms long filaments with Cdc31 and serves as a scaffold for spindle pole body duplication. C-terminal domains interact with other Sfi1 molecules, allowing for anti-parallel dimerization^28^.

Evidence in ciliates shows that Sfi1 and centrin complexes link together to form a dense, fishnet-like mesh.^30,34,35^ The *Paramecium* Sfi1 ortholog, PtCenBP1 is of similar size and structure as *Stentor* Sfi1, encoding a 3.8k -aa protein with a theoretical molecular mass of 460kDa of largely alpha-helical structure with coiled-coiled arrangements and the N and C termini, consistent with a filamentous protein able to form homopolymers^30^. The dem1 mutant, which contains a truncated Pt*CenBP1* is characterized by having sparse ICL meshes and decreased branching^30^, much like what we observe in *Stentor*, Sfi1-1 and Sfi1-2 knockdowns.

*Spirostomum minus* Sfi1 orthologs GSBP1 and GSBP2 are much larger (1458kDa and 2008 kDa) and contain 184 and 267 Sfi1 repeats^34^. These large proteins, along with its centrin family binding partner spasmin, are able to form protein complexes that resemble ropes that are part of a contractile mesh-like fibrillar system. This mesh is organized as numerous parallelograms, much like the *Paramecium* ICL. The parallelograms shrink in the cell’s contracted state, causing the fibrillar bundles to shorten and thicken^34^. In contrast to such regular geometric arrangements, *Stentor* Sfi1 anterior mesh appears with an irregular, banded appearance. Studies have shown that myoneme bundles in *Stentor* do shorten and thicken with contraction^36,37^, however, we have not been able to observe the difference in the mesh in immunofluorescence of extended and contracted cells due to rapid contraction of cells upon fixation. In *Spirostomum minus*, contractility was tested using different extracellular Ca^2+^ concentrations, which induces spontaneous contraction of the cell. With increasing Ca^2+^ concentration, the average time required for cell contraction reduced. In RNAi experiments, it was observed that some RNAi cells did not contract even in elevated, suggesting that structural integrity of the contractile myoneme mesh is essential for calcium-induced contractility. Indeed, disruption of the mesh-like scaffold could be observed in these non-contractile cells. These results closely mirror our findings in *Stentor*, although increasing extracellular Ca^2+^ did not result in spontaneous contraction, and thus other external stimuli were used in our experiments to induce contraction, ensuring that the failure to contract in knockdown cells was not due to insufficient signaling input but rather to defects in the contractile machinery itself.

The greatest difference in *Stentor* Sfi1-centrin scaffolds from these other ciliate systems are the polarized regional differences. This may be due to the contrast in cell shapes. *Spirostomum* adopts a long, cylindrical shape that is tapered on both ends and *Paramecium* are ovoid with a rounded anterior and slightly pointed posterior. These cells contract along their entire body axis, and the geometric mesh of the contractile centrin cytoskeleton in these ciliates is largely uniform across the cell bodies. In contrast, *Stentor* cells are trumpet-shaped, with a flared anterior and a stalk-like tail at its posterior. Although this shape may bear greater resemblance to another contractile ciliate *Vorticella*, the stalk of *Vorticella*, containing the highly contractile spasmoneme, is a separate structure from the zooid or cell body, and contracts like a spring. In *Stentor*, the entire cell contracts, with its posterior being more contractile than its anterior. Another ciliate with a highly polarized cell shape is *Lacrymaria*, a tear-drop shaped predatory ciliate with a highly contractile neck. Like *Stentor*, its centrin cytoskeleton has differences in anterior-posterior patterning^12,52^. In the contractile neck of *Lacrymaria*, centrin forms strand-like fibers, whereas in the cell body, it is distributed in a network. However, in a proteomic analysis, Sfi1 homologs are not found in the *Lacrymari*a^52^.Contraction in this system, likely involves an actin-myosin system^52^. While the Sfi1-centrin complex is a broadly conserved feature of contractile cytoskeletons across ciliates, its structural organization and functional deployment are uniquely adapted to each organism’s cell shape, contractile demands, and spatial patterning, with *Stentor* exhibiting particularly polarized and specialized architecture.

### Sequential scaffolding

Our data suggest that Sfi1 proteins are recruited to the oral primordium in a defined temporal order, supporting the model that the scaffold required for primordium growth is assembled sequentially. Among these, Sfi1-3 appears to be recruited in later stages of regeneration, and its knockdown correspondingly results in minimal or negligible defects in regeneration. This suggests that Sfi1-3 may not be essential for early stages of scaffold formation or that functional redundancy among Sfi1 isoforms allows other Sfi1 proteins to compensate for its absence. In contrast, Sfi1-1 and Sfi1-2, which are expressed earlier during regeneration, may play more central roles in establishing the structural framework of the developing primordium. Their knockdowns lead to more severe phenotypes, implying that the early scaffolds they form are more critical to initiate and sustain proper morphogenesis.

We propose a model in which Sfi1 isoforms operate in a relay-like sequence, with each variant scaffolding centrin assemblies at specific developmental timepoints. For example, Sfi1-1 may scaffold the early primordium, after which it is removed or turned over. Subsequently, Sfi1-2 replaces or supplements this scaffold during an intermediate stage, followed by Sfi1-3 contributing to late-stage organization of the newly formed oral apparatus. We hypothesize that this stepwise scaffold handoff allows for stage specific remodeling of the cytoskeleton, enabling dynamic transitions in structural organization without compromising overall spatial integrity.

This mechanism is reminiscent of T4 bacteriophage assembly, where a temporary internal scaffold is used to shape the prohead (capsid precursor). The protein constituents of the assembly core, p22 and internal proteins, are gene products which are components of the phage head precursor but are not structural components of the finished capsid^58^. During bacteriophage morphogenesis, specific scaffold proteins are recruited in a temporal order, and once their role is fulfilled, they are proteolytically degraded or removed, allowing the structure to advance to the next stage. These scaffolds are essential for proper assembly but are not incorporated into the final mature virion. Similarly, in ribosome biogenesis, pre-rRNA molecules are synthesized and associated with ribosomal proteins and ribosome biogenesis factors^59^. These ribosome biogenesis factors temporarily associate with the pre-ribosomes, and bind in a hierarchical and sequential manner, where association with earlier factors is required for the association of later factors. This stepwise assembly drives the maturation of the ribosomal subunits, without being a part of the mature ribosome^59,60^. In *Stentor*, different Sfi1 proteins may act as transient guides, sequentially shaping the growing primordium, but not necessarily remaining part of the final, completed oral apparatus. This sequential assembly ensures that correct proteins are localized at the correct time and place, contributing to the efficient and accurate assembly of complex structures.

### Role of lateral filaments as potential circumferential positional cue

Classical studies on insect limb development have shown that when regions containing different positional cues are placed in proximity, it can trigger the formation of new organizing centers^61^, development of ectopic structures, or in some cases intercalation of limb regions in inverted orientations^62^. Such observations point to mechanism for comparing positional information in neighboring regions so as to ensure continuity of form. Intracellular juxtaposition can likewise play a pivotal role in establishing defined positions in cellular patterning with *Stentor* providing a classic example. In *Stentor*, grafting pieces of the cortex with different stripe widths not containing an oral primordium site is sufficient to induce the formation of a new primordium^63^. This observation emphasizes the critical role of the juxtaposition of cortical regions with differing stripe widths appearing as a key organizing factor, with circumferential cues helping to establish a radial pattern that governs the location of oral primordium formation. However, the molecular basis for such circumferential positional information in *Stentor* are completely unknown.

Whatever the circumferential position information is based on, it correlates with stripe width which reflects the spacing between adjacent microtubule bundles (km fibers). Because the centrin cables are aligned with the microtubule bundles, stripe width thus also correlates with the spacing between longitudinal centrin cables. The spacing of the cables decreases as one moves counterclockwise around the cell when viewed from its anterior pole, terminating at an intersection of narrow and wide-spaced cables. This “locus of stripe contrast” defines the oral primordium site. When stripe contrast regions are created by surgical manipulation in different regions of the cell, they are sufficient to produce a new oral primordium, strongly suggesting that all cortical stripes contain some form of positional information.

Given that Sfi1 bands span the width between adjacent ciliary rows and show distinct expression patterns along the anterior-posterior axis, we propose that the length of these filaments may encode positional information. This model raises two questions. First, how are differences in the length of the Sfi1 bands read out to determine the site of primordium development, and second, how is a smoothly varying gradient of band lengths created. We currently have no answer to the first question, but we can propose a hypothesis for the second. It is known that the number of stripes increases with the size of the cell. Stripe multiplication can occur by splitting or “branching” where the widest pigment stripes begin to split when a new clear stripe of ciliary rows and fibrous structures emerges within the pigmented stripe. This growth process follows a spiral trajectory, with new short stripes added anteriorly and older stripes extending posteriorly, eventually reaching the posterior pole^6^. This would indicate that the newest ciliary rows are more tightly packed and shorter in length along the longitudinal axis of the cell, compared to older stripes which are longer and more widely spaced. Sfi1 bands might thus reflect the developmental age of the ciliary stripes or serve as landmarks that may help establish circumferential identity across the cortex. If existing Sfi1 bands undergo gradual elongation over successive rounds of cell division, for example by incorporation of more Sfi1-centrin superhelical filaments, this would produce the observed circumferential gradient in filament length.

The potential role of Sfi1 bands as structural and spatial reference points is consistent with a model in which Stentor uses a cytoskeletal scaffold not just for mechanical support but for spatial patterning in which it integrates both longitudinal and circumferential positional cues to coordinate morphogenesis.

## Conclusion

Our Sfi1 knockdown experiments demonstrate that loss of Sfi1 impairs not only oral apparatus regeneration but also whole-cell contractility, providing direct functional evidence that Sfi1 is required for both morphogenetic and mechanical processes. These findings suggest that in *Stentor*, Sfi1 links the molecular architecture of the cortex to the mechanical behavior of the entire cell, a role that likely evolved to support the unique demands of such a large, single-celled organism. Together, our findings suggest that spatial patterns at the sub-cellular level can be generated by spatiotemporal variation of scaffolding proteins that, in turn, organize structural proteins into distinct structures at the appropriate time and place during cellular development.

## Materials and methods

### Stentor culturing and media

*Stentor coeruleus* cells were obtained from Carolina Biological Supply and grown in the dark, at room temperature in the lab. Cells are cultured in pasteurized spring water (Carolina Biological Supply). Cells are fed every 4-5 days with *Chlamydomonas reinhardtii* grown on TAP agar plates and resuspended in 15mL pasteurized spring water, with each 2qt culture being fed 5mL.

### Molecular Phylogenetics

For the phylogenetic characterization of *Stentor* candidate centrin and Sfi1 sequences, we gathered amino acid sequences of human, *Saccharomyces cereviasiae*, *Tetrahymena thermophila*, *Blepharisma stoltei* or *Paramecium tetraurelia* homologs using online BLASTP^64^ and human centrin 2/ Sfi1 as queries. We aligned them with *Stentor* sequences using MUSCLE (v3.8.425) and corrected the alignments manually by deleting gap positions and sequenced that could not be aligned properly. We used IQ-TREE2 (v2.2.0.3) with the Le-Gascuel substitution matrix and gamma distribution of among site rate-variation^65^ and tested for robustness of the topologies by running 1000 ultrafast bootstraps^66^.

### Sfi1 Analysis

Sfi1 homologs were identified in *Stentor* using BLAST^64^, RADAR (Rapid Automatic Detection and Alignment of Repeats)^43^ and ScanProsite^44^. Potential centrin binding sites were found using RADAR with repeats ending in W[KR]. Sequence logos generated from SteCoe_25973, SteCoe_10021, and SteCoe_17904 using WebLogo (Berkeley) showed the same sequence. Phylogenetic trees were generated using the Maximum Likelihood Method^67^.

### Alphafold3 structure predictions

The seven predicted Sfi1 repeats used in Alphafold predictions in Figure 2E and Figure 2F spans the amino acids in the positions 753-1293 of SteCoe_17904. The sequence of SteCoe_3383 was used for *Stentor* centrin 3.

### Antibody production

Pre-immune bleeds from rabbits were screened to avoid using rabbits previously infected with ciliated parasites that may produce antibodies that react with *Stentor* proteins. Screening was done by incubating western blots of *Stentor* proteins with pre-immune rabbit serum at a 1:500 dilution, then staining with secondary antibodies. Custom anti-Sfi1-1, anti-Sfi1-2, and anti-Sfi1-3 antibodies were generated and purified (Fortis Labs, Richardson TX) using the peptides RKKEDKNQVRKTP, MEGSKIPTLLED, ENIIRRTMRDADQR respectively. Antibodies were validated using a peptide block and immunofluorescence. Peptides were incubated at a 1:10 Ab:peptide ratio and incubated for 2 hours at room temperature.

### Western blots

Lysate samples were prepared using 4X NuPage LDS sample buffer and run on Invitrogen Bolt bis-Tris Mini Protein Gels run in a Mini Gel Tank (Fisher Scientific A25977) and transferred using the accompanying blotting module. HiMark^TM^ pre-stained protein standard was used for Sfi1 western blots with high molecular weight, while PageRuler Plus Prestained protein ladder used in centrin western blots. Membranes were blocked for 1hr at RT with 3% BSA in PBST. Primary antibodies were diluted at 1:200 and incubated at 4C overnight. Secondary HRP antibodies were diluted at 1:1000 and incubated for 1hr at RT. Super Signal West Pico Chemi Substrate used for protein detection.

### RNAi constructs

Sfi1 genes were amplified using PCR from *Stentor* genomic DNA isolated using a DNeasy Blood and Tissue Kit: Animal Blood Spin-Column Protocol (Qiagen, Germantown, MD). Ligation independent cloning was used to insert gene fragments into a pPR-T4p plasmid between two T7 promoters, and the vectors were transformed into Dh5a *E. coli* cells. Primer sequences and targeted regions are indicated in Supplementary Table 1. For negative controls, either empty pPR-T4p vectors containing no insertion or constructs cloned with the LF4 gene, which is involved in cilia length control but otherwise does not affect function or morphogenesis, were used.

### RNA interference

RNAi was either performed by feeding^68^ or microinjection. Feeding constructs were transformed into HT115 *E. coli* and grown to OD 0.4-0.6, then induced with 1mM isopropyl-β-d-thiogalactopyranoside (IPTG) and incubated shaking overnight at room temperature. *Stentor* were fed daily with resuspended bacteria pellets for 7 days. Wells were cleaned and replaced with fresh Carolina Spring Water every other day. Microinjection protocol was adapted from Slabodnick, 2023 (unpublished). Briefly, 70-100 cells were placed in chambered slides containing 2mL 1.5% methylcellulose. Cells were allowed to acclimate and fully extend. Drummond Borosilicate glass capillaries (3-000-203-G/X) were siliconized using Sigma Aldrich SigmaCote (SL2) and allowed to air dry. Capillaries were pulled using a Sutter micropipette puller (P-97) and needles were cut with a razor blade. RNA was loaded into needles using Microloader pipette tips (Calibre Scientific). RNA was synthesized using NEB HiScribe T7 High Yield RNA Synthesis kit (E2040S) using linearized pPR-T4p vector with cloned constructs as the template. RNA was concentrated and cleaned using NEB Monarch Spin RNA Cleanup Kit 500ug (T2050L) and microinjected at 800-2000ng/uL using a Drummond Nanoject II microinjector.

### Regeneration Assay

Sucrose shock regeneration assay is adapted from Lin (2022). Briefly, *Stentor* were concentrated in 1mL in a 50mL conical tube. 1mL 30% sucrose (Sigma Aldrich S7903) was added for a final concentration of 15% and gently mixed. Cells were inspected visually for 1-2min until the membranellar bands had fallen off. Cells were washed three times in pasteurized spring water (Carolina Biological Supply) to remove sucrose and visually staged at 4, 6, 8, and 24 hours post sucrose shock. For easier staging, 20-30 cells were placed in a 9mm spacer (Grace BioLabs) on a microscope slide, sealed with a 18x18 glass coverslip and observed under a Zeiss AxioZoom V16 microscope.

For regeneration after bisection, cells were cut in half as previously described^69^. Cells were placed in 0.25% methylcellulose on parafilm and cut with a glass needle. Posterior and anterior halves were separated and washed three times to remove methylcellulose.

### Immunofluorescence and Expansion

Cells were fixed in freezing methanol (Sigma Aldrich 67-56-1) overnight and rehydrated in methanol in 0.5x PBS for 20 minutes. Cells were washed 3 times with 0.1% TBS-Triton X and blocked in antibody dilution buffer containing 2.5% BSA (Sigma Aldrich A030), 0.5% cold water fish gelatin (Sigma Aldrich G041), 0.1% Triton X in PBS overnight at RT. Primary antibodies were added using the following dilutions: Centrin 20H5 (Sigma Aldrich 04-1624) 1:100, Sfi1-1-3 1:100, and Actub 6-11B-1 (Sigma Aldrich T7451) 1:1000 and incubated for 1 hour at RT and washed 3x with TBS-Tx. Secondary antibodies goat anti-mouse 488 (Invitrogen A-11001), goat anti-mouse 594 (Invitrogen), goat anti-rabbit 488 (Invitrogen) and goat anti-rabbit 594 (Invitrogen) were used at a 1:200 dilution and incubated for 1 hour at RT. Cells were then washed 3x with TBS-Tx. Cells were mounted in Prolonged Glass Antifade (Invitrogen P36980) and placed inside a 9mm SecureSeal spacer (Grace BioLabs) adhered to a microscope slide and sealed with a 18x18mm glass coverslip. For expansion samples, cells were fixed and stained as described above. A modified Magnify^47^ expansion protocol was used for expansion. Briefly, fixed cells were mounted on poly-lysine coated 8mm round coverslips (Thomas Scientific 127H20) and embedded in the Magnify gel monomer solution overnight at 37C in a humidified chamber. Gels were carefully removed and placed in an Eppendorf tube with 1mL Magnify homogenization buffer and incubated at 80C for 24 hours. Gels were washed in 1xPBS. Gels were stained again using 2x the antibody concentration used in pre-gelation samples diluted in 2% BSA in PBS at RT for >2 hours. After washing 3x 10min each in 1xPBS, DyLight 594 NHS Ester (Thermo Scientific PI46413) was added at 20ug/mL in addition to any other secondary antibodies used. Gels were washed 3x10min in 1xPBS and expanded twice for 30min each in ddH20, then expanded once more overnight. Gels were imaged in Ibidi 2-well slides and imaged using confocal microscopy.

### Imaging

Brightfield imaging was done using Zeiss AxioZoom V16 equipped with a Nikon Rebel T3i SLR Camera. Confocal microscopy was done using CSU-WI Spinning disk equipped with an Andor Zyla sCMOS camera. High speed imaging was done using Vision Research v7.3 or v341 high speed cameras.

### Contraction Assays

Pipetting Assay: Fully extended free-floating or anchored cells were pipetted into a P20 pipette tip, which induced contraction in control cells. Each cell was not assayed more than once to prevent habituation.

High-Speed Contraction Assay: Individual cells were imaged in a 9x9 SecureSeal spacer (GraceBios) adhered onto a microscope slide and sealed with a glass coverslip. A mechanical force onto the side of the slide was used to induce contraction of fully extended cells.

Electrical Pulse Assay: *Stentor* cells were stimulated to contract using an electrical pulse. A 555 timer-based one-shot pulse generator (a kind gift from Joseph Lannan) with a potentiometer (Bourns Inc 2590S-2-103L) was attached to glass slides with flat pieces of metal obtained by breaking a double-edged razor blade, on the longitudinal sides functioning as both electrodes and spacers. These pieces of metal and vacuum grease created a chamber in which 20-50 cells in 1.5% methylcellulose were placed and gently compressed with a coverslip. The methylcellulose along with the compression from the coverslip helps to promote full extension of the cells. Images before and immediately after the electrical pulse were taken using a USB microscope (Celestron Handheld Digital Microscope Pro).

## Supporting information

alphafold files supplement 1

alphafold files supplement 2

alphafold files supplement 3

Supplemental videos of cell contraction events

## Acknowledgements

This material is based upon work supported by the National Science Foundation Graduate Research Fellowship Program under Grant No. 2038436. Any opinions, findings, and conclusions or recommendations expressed in this material are those of the author(s) and do not necessarily reflect the views of the National Science Foundation. This work was supported by NIH grant R35 GM130327. We would like to thank Kaustubh Ramachandran for helpful discussions about centrin phylogeny, Mary Elting, Joe Lannan, and Jerry Honts for discussions about Sfi1 in other contractile ciliates. We thank Rajorshi Paul and Ambika Nadkarni for help with high-speed imaging, and we thank members of the Marshall lab for helpful discussions.

---

**Supplementary Figure 1:**
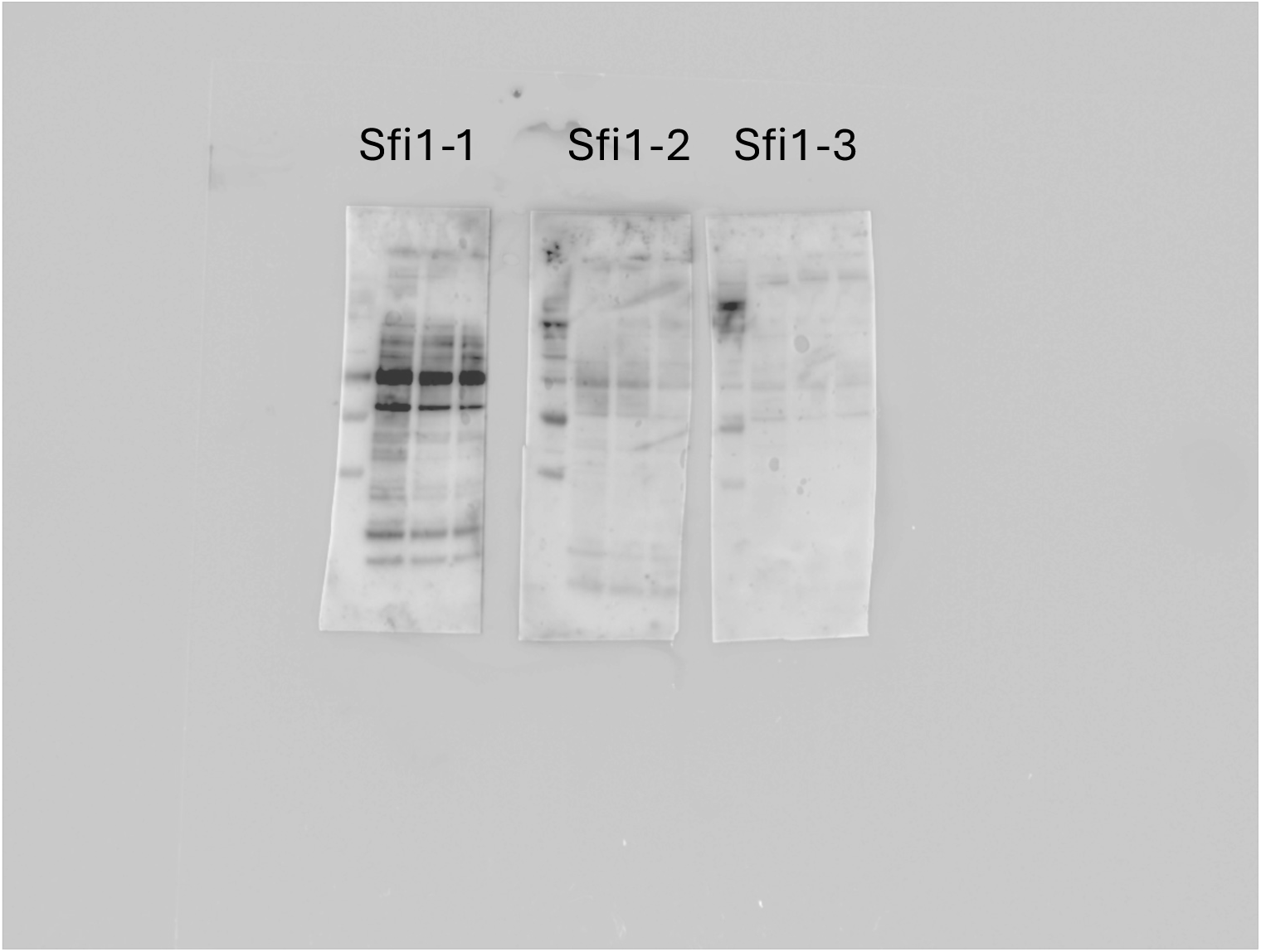
Sfi1 Western blots. Separate blots for Sfi1-1, Sfi1-2, and Sfi1-3 with HiMark ladder on the left and three lanes of lysate blot with the corresponding antibody.

**Supplementary Figure 2:**
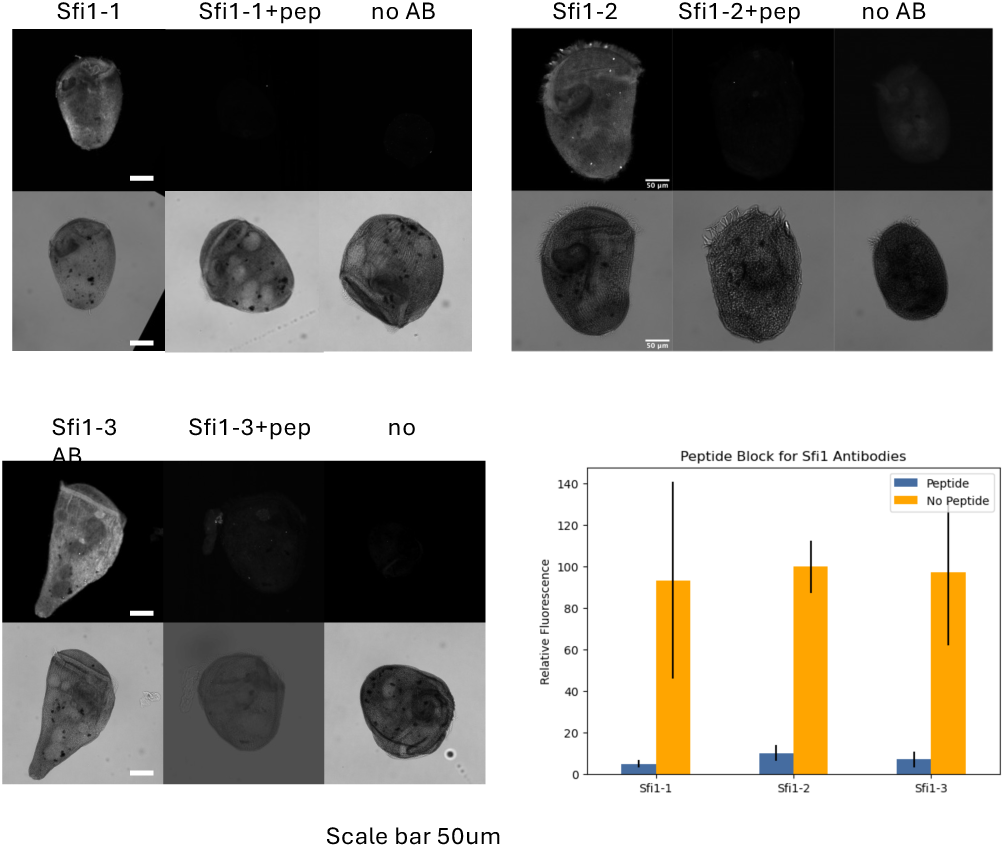
Sfi1 antibody validation with peptide block. Immunofluorescence with peptide incubated antibodies and no primary controls reveals little to no signal compared to the antibody without a peptide block. Brightfield images of cells shown on bottom panel.

**Supplementary Figure 3:**
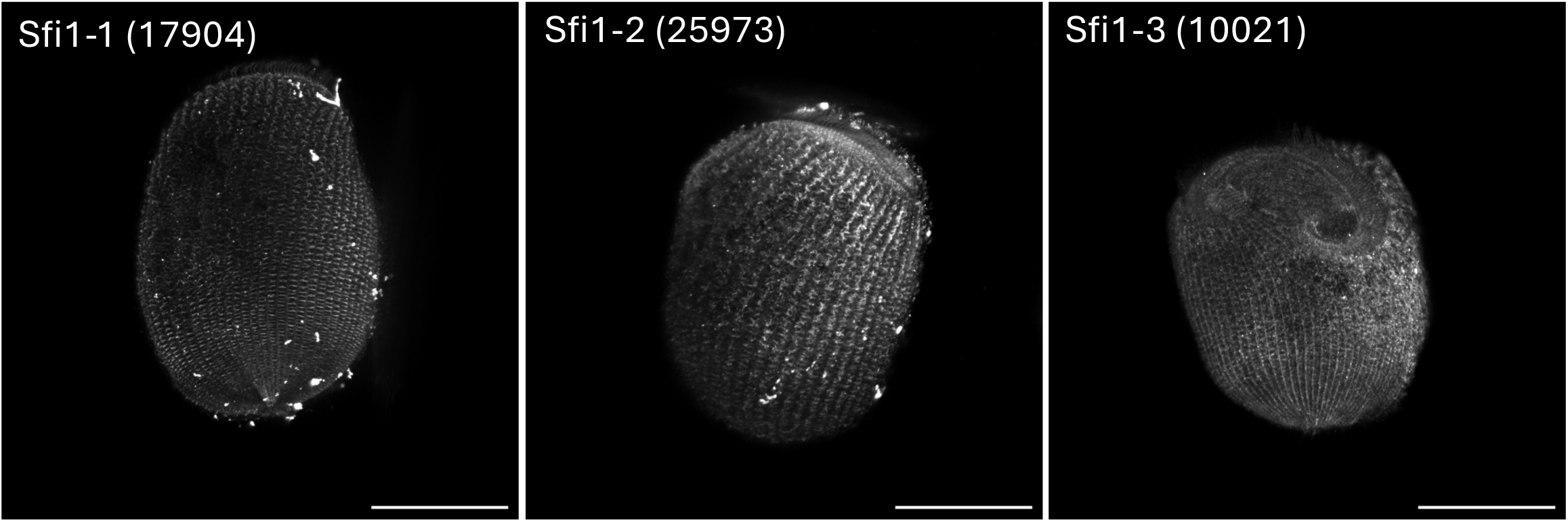
Representative staining for each Sfi1 antibody. Representative maxium intensity projection confocal z-stacks of Sfi1-1, Sfi1-2, and Sfi1-3 antibodies all shown similar localization patterns.

**Supplementary Figure 4:**
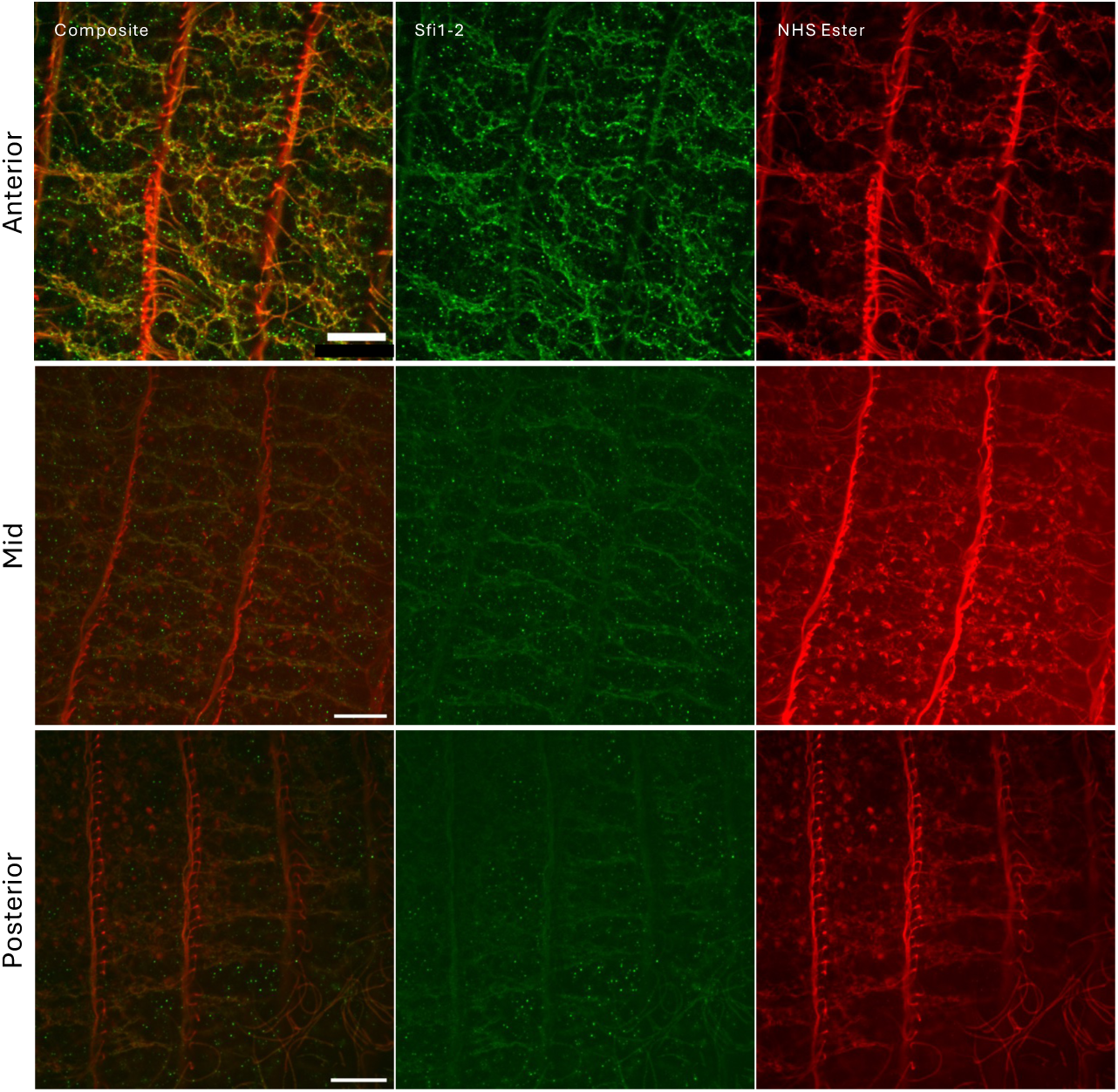
NHS Ester labels Sfi1 structures. Expansion microscopy of Sfi1 structures labeled with Sfi1-2 and NHS-ester.

**Supplementary Figure 5:**
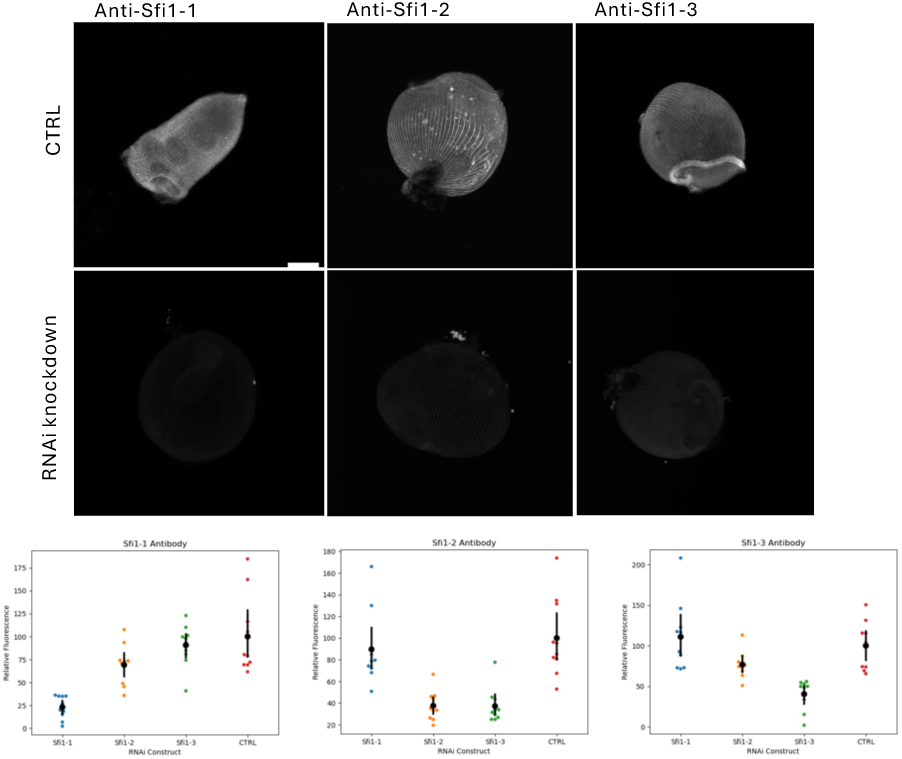
Antibody validation of Sfi1 knockdowns. Sfi1s are knocked down using RNAi and stained with the corresponding antibody, revealing little to no signal in the knockdowns. Cross antibody/knockdown staining shows some overlap between Sfi1-2 and Sfi1-3.

**Supplementary Figure 6:**
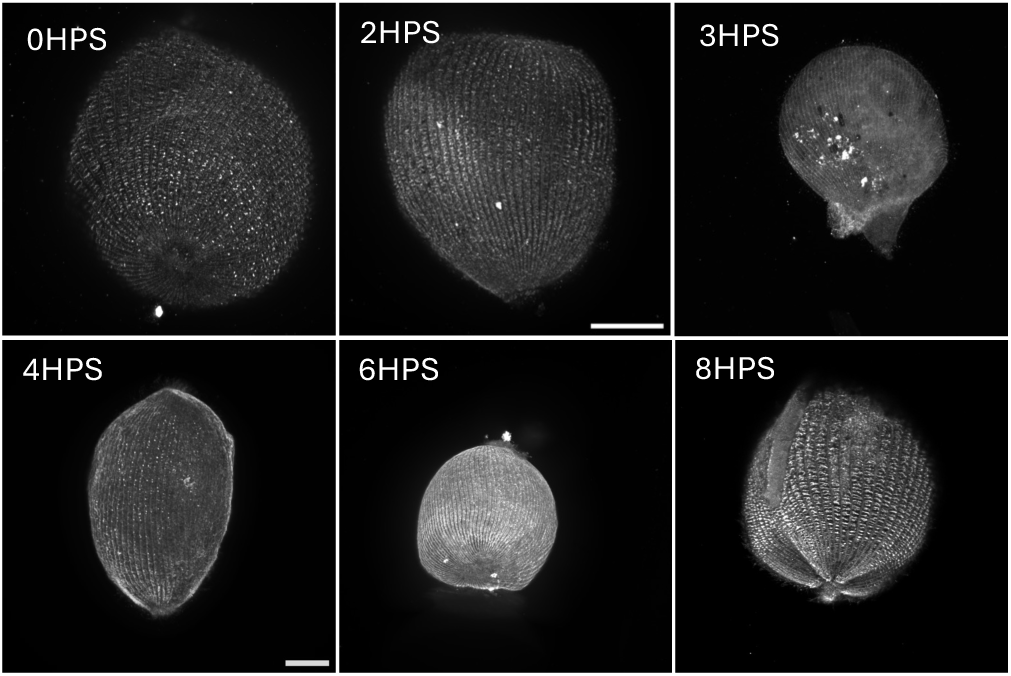
Re-establishment of Sfi1 bands during regeneration. Representative maximum intensity projection confocal z-stacks of Stentor cells at timepoints after removal of its oral apparatus stained with anti-Sfi1-2. Immediately after sucrose shock, the Sfi1 bands appear faint and more punctate at 2HPS. By 3HPS, the bands are not visible, leaving only the Sfi1 in the cortical fibers. Sfi1 puncta begin to appear at 4HPS and fully re-establish at 8HPS.

**Supplementary Videos:**
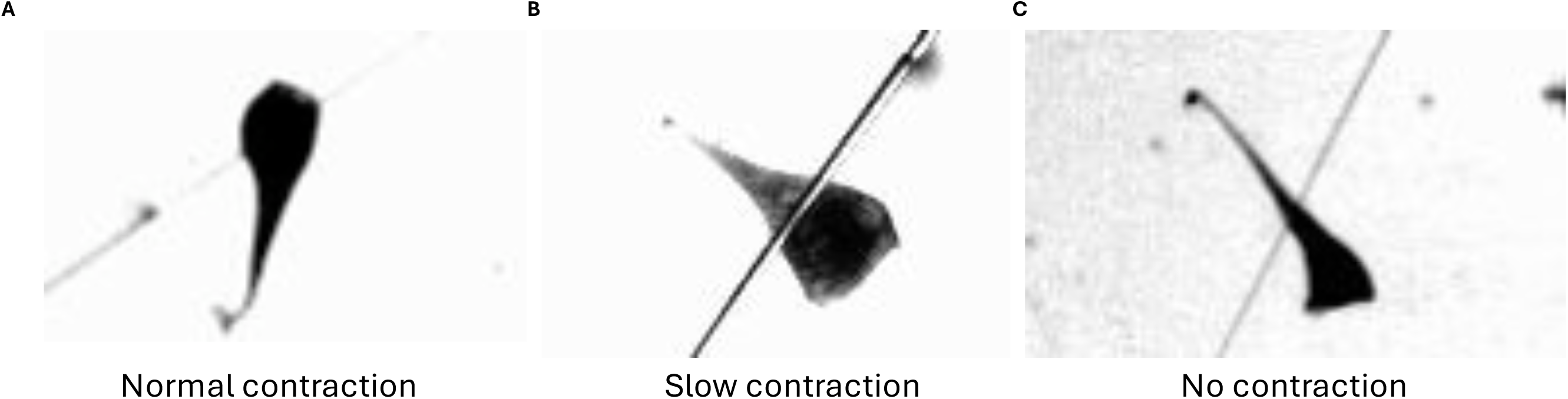
Representative videos of normal contracting, slow contracting, and no contraction cells in response to a mechanical stimulus.

**Supplementary Table 1:**
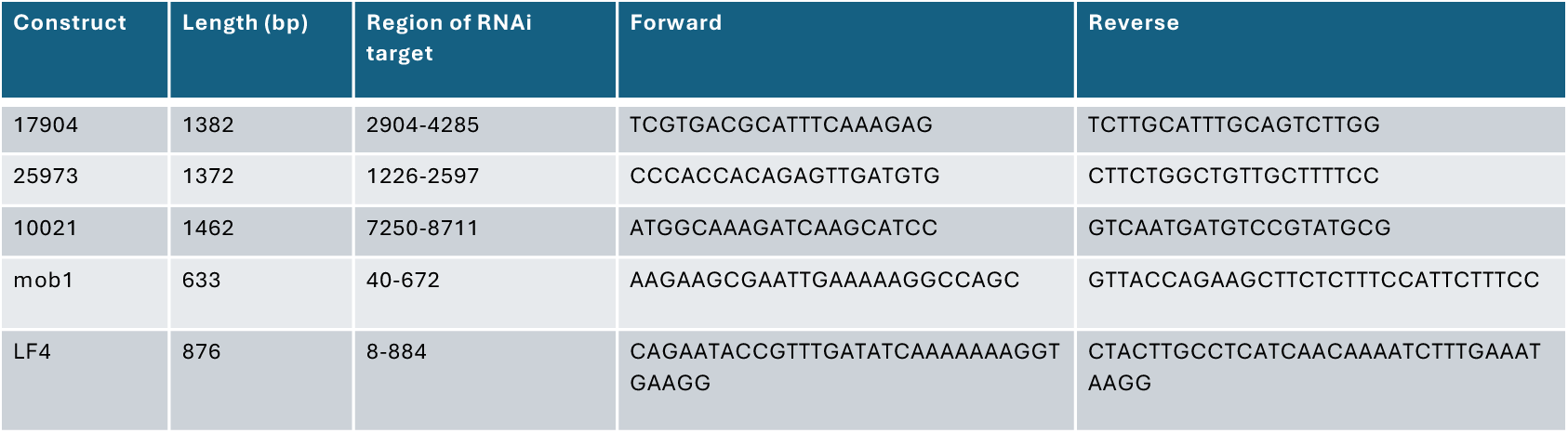
RNAi contruct primers. Primers used to generate Sfi1 knockdown constructs and controls. Mob1 and LF4 primers are from Slabodnick (2014).

