## Supplemental videos of cell contraction events for "Spatiotemporal control of cortical centrin patterning by regionalized Sfi1 family scaffolding proteins in *Stentor coeruleus*"

### Slide 1
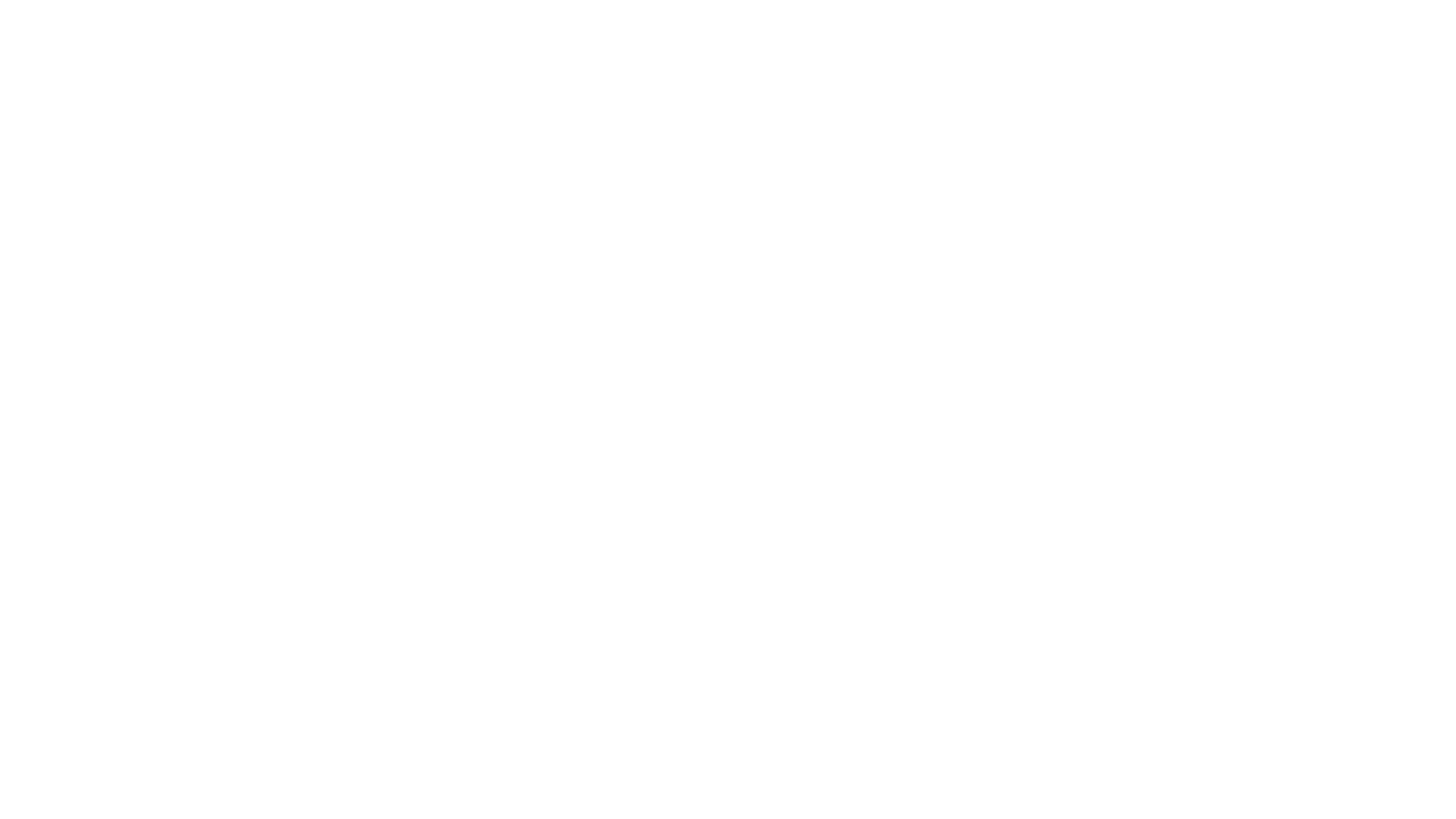

#

### Slide 2
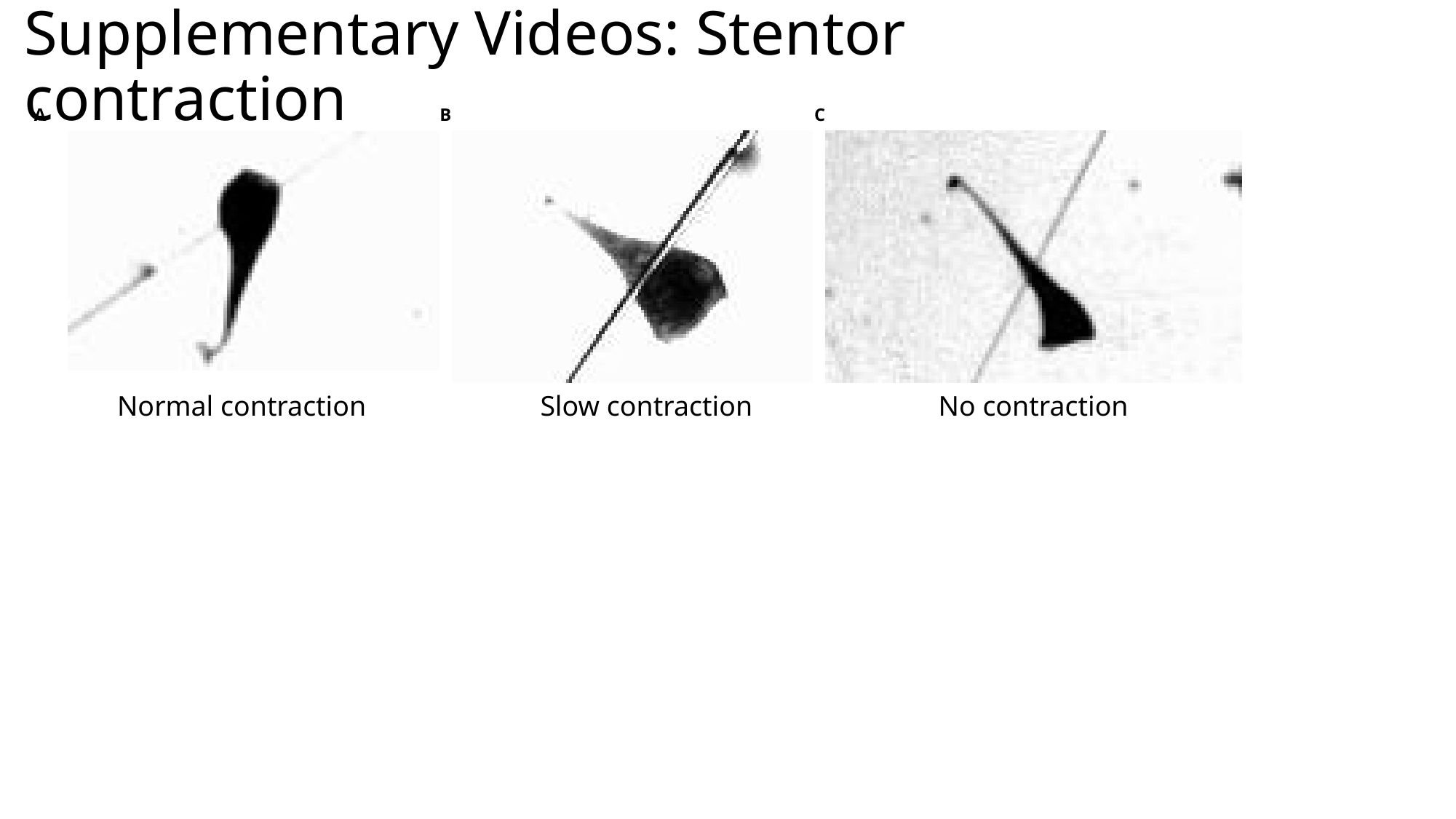

#### Supplementary Videos: Stentor contraction
B
C
A
Normal contraction
Slow contraction
No contraction
